# Multiple brace root phenotypes promote anchorage and limit root lodging in maize

**DOI:** 10.1101/2021.05.12.443923

**Authors:** Ashley N. Hostetler, Lindsay Erndwein, Jonathan W. Reneau, Adam Stager, Herbert G. Tanner, Douglas Cook, Erin E. Sparks

## Abstract

Plant mechanical failure (lodging) causes global yield losses of 7-66% in cereal crops. We have previously shown that the above-ground nodal roots (brace roots) in maize are critical for anchorage. However, it is unknown how brace root phenotypes vary across genotypes and the functional consequence of this variation. This study quantifies the contribution of brace roots to anchorage, brace root traits, plant height, and root lodging susceptibility in 52 maize inbred lines. We show that the contribution of brace roots to anchorage and root lodging susceptibility varies among genotypes and this contribution can be explained by plant architectural variation. Additionally, supervised machine learning models were developed and show that multiple plant architectural phenotypes can predict the contribution of brace roots to anchorage and root lodging susceptibility. Together these data define the plant architectures that are important in lodging resistance and show that the contribution of brace roots to anchorage is a good proxy for root lodging susceptibility.

## Introduction

A changing global climate is increasing the severity and prevalence of storm systems, which threatens crop production. Crop failure from mechanical stress is called lodging and is defined as the displacement of plants from vertical (Rajkumara, 2008). In cereal crops, lodging is reported to cause between 7-66% yield loss (Carter and Hudelson, 1988; Flint□Garcia et al., 2003; Rajkumara, 2008; Tirado et al., 2020) and can be attributed to stalk and/or root failure (Berry et al., 2004; Rajkumara, 2008). Stalk lodging occurs at late growth stages when internodes buckle below the ear, whereas root lodging can occur at any growth stage when the root system fails by uprooting and/or breaking (Berry et al., 2004; Rajkumara, 2008; Erndwein et al., 2020; Hostetler et al., 2021). The development of lodging-resistant crops relies on a detailed understanding of the underlying factors that contribute to lodging susceptibility. However, defining these factors has been limited by the reliance on unpredictable weather patterns to induce natural lodging events. To overcome this limitation, measures of plant anchorage have been developed as a proxy for root lodging (Erndwein et al., 2020). Using both natural lodging events and measures of plant anchorage, several factors that impact lodging have been defined, and include field management practices, plant stage at lodging, plant genotype, and plant architecture (e.g. root system architecture and plant height) (Carter and Hudelson, 1988; Berry et al., 2004; Rajkumara, 2008).

In maize, the size and extent of the root system has been linked to root lodging resistance and anchorage (Thompson, 1972; Sanguineti et al., 1998; Liu et al., 2012; Farkhari et al., 2013; Sharma and Carena, 2016; Shi et al., 2019). Within the root system, the presence of above-ground nodal roots (called brace roots) is consistently associated with the ability to resist root lodging (Hoppe et al., 1986; Carter and Hudelson, 1988; Stamp and Kiel, 1992; Liu et al., 2012; Sharma and Carena, 2016; Shi et al., 2019; Blizard and Sparks, 2020; Hostetler et al., 2021). However, beyond quantifying the presence of brace roots, there have been limited studies that quantify other root phenotypes that may contribute to root lodging resistance and anchorage. A handful of studies have identified the number of brace root whorls in the ground (Liu et al., 2012; Sharma and Carena, 2016; Shi et al., 2019; Reneau et al., 2020), the number of brace roots in a whorl (Liu et al., 2012; Sharma and Carena, 2016), the brace root diameter (Liu et al., 2012; Sharma and Carena, 2016), and the brace root spread width (Sharma and Carena, 2016) as important phenotypes for root lodging resistance. In addition to root phenotypes, plant height is a known factor in lodging susceptibility for many crops (Rajkumara, 2008). Yet in maize the link between plant height and root lodging is unclear. One study showed no correlation between plant height and root lodging in a natural root lodging event (Sharma and Carena, 2016), whereas another study showed plant height as a secondary factor in root lodging models, which were validated by natural root lodging events (Brune et al., 2018). The wide-spread interpretation of previous results has been limited by a small number of genotypes per study, disparate phenotypes analyzed in each study, and a primary reliance on univariate correlations. Therefore, a detailed assessment of phenotypes at the population level is required to define how plant architectures contribute to root lodging resistance and anchorage.

In this study, the brace root contribution to anchorage, brace root phenotypes, and plant height were assessed in a population of 52 maize inbred lines. Additionally, root lodging susceptibility was quantified after a natural storm event within the same population. Within this population there was substantial variation in the brace root contribution to anchorage, plant architectural traits (brace root phenotypes and plant height), and root lodging susceptibility. Consistent with previous results, individual phenotype correlations with the brace root contribution to anchorage and root lodging were moderate. However, when using supervised classification models, the brace root contribution to stalk anchorage and root lodging were classified with high accuracy. Multiple plant phenotypes drove the decision tree, thus highlighting the importance of multi-trait analyses for identifying plant architectures that impact lodging susceptibility. Further, this study demonstrates a linkage between plant height and brace root traits, which have opposing effects on lodging susceptibility.

## Materials and Methods

### Plant Material

A maize (*Zea mays*) population, consisting of 52 inbred genotypes with genetic and phenotypic diversity (Table S1), was grown in the summers of 2018, 2019, and 2020 at our Newark, DE field site (39° 40’ N and 75° 45’ W) with overhead irrigation. Each year, 2-4 replicate plots (12’ single row) of eighteen seeds per plot were planted in a randomized design. Seeds were planted on 5/30/2018, 5/22/2019, and 06/01/2020. Monthly temperature averages, rainfall estimates, and wind averages can be found in Table S2. Additional weather data can be found at the Delaware Environmental Observing System (http://www.deos.udel.edu) and selecting the Newark, DE-Ag Farm station. Pre-emergence (Lexar at 3.5 quart per acre and Simazine at 1.2 quart per acre) and post-emergence (Accent at 0.67 ounces per acre) herbicides were applied. Soil insecticide COUNTER 20G was applied at 5.5 lbs per acre at planting. Ammonium Sulfate 21-0-0 was applied at 90 lbs per acre at planting, and 30% Urea Ammonium Nitrate (UAN) was applied at 40 gallons per acre approximately 1-month after planting (at knee high).

### Field Based Measurements of Plant Biomechanics

Non-destructive, field-based measurements of plant biomechanics were collected with the <u>D</u>evice for <u>A</u>ssessing <u>R</u>esistance to <u>L</u>odging <u>IN G</u>rains (DARLING) (Cook et al., 2019) as described in Reneau et al., (2020). In 2018, 3-5 plants per plot were tested at 144-165 DAP. In 2019, all viable plants were tested at 118-129 DAP. Plants were tested after dry down to avoid daytime variation as previously reported (Reneau et al., 2020). Plants were excluded from testing if their root systems were overlapping with neighboring plants, there was disease or pest damage, or plants were damaged during weed control measures. For each test, the pivot point of the device was aligned with the base of the plant, and the load cell was placed in contact with the stalk at 0.64 m. Three cycles of deflection, each displacing the stalk approximately 15-degrees before returning to 0-degrees, were applied to each plant. Output of each test is a .csv file of force-rotation data. Rotation, measured from an inertial measurement unit (IMU), was converted from degrees to radians with the following equation:

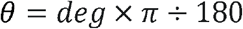

where *π* is taken as 3.1415. The rotation in radians was then converted into deflection (*δ*) with the following equation:

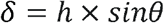

where *h* is the height of the applied load (0.64 m) and *θ* is the rotational angle (radians). The slope of the Force-Deflection data (N/m) was determined by fitting a line to the entire dataset (loading and unloading data for all three cycles). We previously determined that this data extraction method is reliable and comparable to extracting the slope of just the loading data (Reneau et al., 2020).

To determine the relative contribution of brace roots to anchorage, all brace roots entering the ground were manually excised with pruning shears and Force-Deflection was remeasured in the absence of brace roots. The relative contribution of brace roots was calculated as the ratio between the slope of the Force-Deflection curve with all brace roots removed (None), and the slope of the Force-Deflection curve with all brace roots intact (All; Ratio=None/All). We have previously shown that repeat testing does not alter the slope of the Force-Deflection curve and any changes in Force-Deflection are due to the removal of brace roots (Reneau et al., 2020). Thus, a lower ratio is indicative of a greater contribution of brace roots to anchorage.

### Plant Phenotyping

In 2019, brace root images were manually acquired with a digital camera. At 107 DAP, fields were manually cleared of any weeds or debris, including lower stem leaves. A 0.5-inch label, which included plot number, plant number, and an A or B designation to identify each side of the plant (Plot_Plant_A or Plot_Plant_B), was then secured to each stem with clear tape. Images were collected from both sides (side A or side B) of 992 plants representing the 52 inbred genotypes. In 2020, brace root images were collected with a ground-based <u>B</u>race <u>R</u>oot phenotyping r<u>obot</u> (BRobot). BRobot is a modified Superdroid LT2 Tracked ATR Robot Platform with a custom controller and side mounted FLIR 3.2 MP Color Blackfly camera (Figure S1). Plots were marked with an orange stake and individual plants were marked with a yellow popsicle stick for reference. Only the number of brace root whorls that entered the ground was quantified from these images.

For the 2019 data analysis, we developed a semi-automated root tagging graphical user interface (GUI) to optimize image processing (Figure S2). This new workflow enables annotation and analysis on the scale of 1-2 minutes per image. The height of the tag adhered to each plant was used as a scale (pixels per inch) and phenotypes were measured for only brace root whorls that entered the ground. These phenotypes are measured sequentially as 1) number of brace roots per whorl (where whorl 1 is closest to the ground), 2) single outermost brace root width for the highest whorl in the ground, 3) stalk width, and 4) a right triangle on each side of the plant from the outermost root from highest whorl in the ground. The right triangle is used to extract the height of the whorl on the stalk (*a* leg of a right triangle), the distance from the stalk-to-root grounding (*c* leg of a right triangle), and the root angle (B angle of the right triangle). The brace root spread width was calculated as the sum of the stalk width and the *b* leg from the triangle on both sides of the plant. The root tagging workflow extracts pixel information and a post processing script converts pixels to inches or degrees. Inches were converted into centimeters for data analysis. Left and right (A and B) phenotyping outputs were compared with a Pearson correlation analysis in R ver. 4.0.2 (R Core Team, 2013) with the *corrplot* package ver. 0.84 (Wei and Simko, 2017). A high correlation (r range 0.55-0.79) between A and B images indicates that the root tagging workflow has high precision (Figure S3), therefore measurements were summed (root number per whorl) or averaged (all other phenotypes) to provide a per-plant brace root phenotypic profile and reduce the effect of phenotyping error. The number of brace root whorls that entered the ground and plant height were manually quantified. Plant height was measured during reproductive development and defined as the distance from the base of the plant to the top of the tassel.

### Lodging Assessment

On August 4^th^, 2020, approximately 65 days after planting (DAP), Tropical Storm Isaias, originally a destructive Category 1 hurricane, caused extensive root lodging. The day after the tropical storm, root lodging data was recorded. For each plot, the total number of plants and total number of root lodged plants were recorded. The percent of root lodging per plot was calculated from these data (number of plants lodged/total number of plants x 100) and was used as a metric for root lodging susceptibility.

### Genotype Relatedness

The phylogenetic relationship among genotypes was determined using PAUP* version 4.0a169 and SVDquartets (Chifman and Kubatko, 2014, 2015; Wascher and Kubatko, 2021). A species tree was generated with exon data, which was downloaded for 41 of the 52 inbred genotypes from the SNPVersity database on MaizeGDB. The exon data downloaded was aligned to the B73 reference genome ver. 2. In instances where a single genotype was represented more than once, all data sets were retained. When the genotype data clustered together, a taxon-partition was used to assign both sets of sequence data to a terminal branch. When genotype data did not cluster, both sets of sequence data were retained as individual branches. The phylogenetic relationships can be visualized in Figure S4.

### Heritability Calculations

Broad-sense heritability (*H*^*2*^) was calculated in R ver. 4.0.2 (R Core Team, 2013) with the following equation:

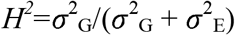

where *σ*^2^_G_ is the genotypic variance and *σ*^2^_E_ is the experimental error variance (Holland et al., 2002). *H*^*2*^ was calculated for the None/All ratio, the architectural phenotypes (root tagging GUI-extracted phenotypes, the manual quantification of the number of brace root whorls in the ground, and plant height), and the root lodging susceptibility data to determine the proportion of phenotypic variance that is due to the underlying genetics.

### Statistical Analyses

All statistical analyses were performed using R ver. 4.0.2 (R Core Team, 2013). An analysis of variance (ANOVA) was performed to determine if there were significant differences among genotypes for each of the architectural traits. Further, a two-way ANOVA was performed to determine the genotype and year effect on the None/All ratio. Prior to running each ANOVA, a Shapiro-Wilk test was used to determine if data was normally distributed. If data deviated from a normal distribution (p<0.05), a Tukey’s Ladder of Powers from the *rcompanion* package ver. 2.3.26 (Mangiafico, 2021) was used to transform the data. Following each ANOVA, if a significant difference was observed, a post-hoc Tukey Honest Significant Difference (HSD) test was performed to test all pairwise comparisons. A principal component analysis (PCA) was used to identify phenotypes that explained the most variability among genotypes. The PCA was visualized with the *factoextra* package ver. 1.0.7 (Kassambara and Mundt, 2020) in R. Pearson correlation analyses were run on: 1) genotypic averages for architectural phenotypes and the None/All ratio for within-year data, 2) genotypic averages for architectural phenotypes and the None/All ratio for multi-year data, and 3) genotypic averages for architectural phenotypes, the None/All ratio, and root lodging susceptibility, with the *corrplot* package ver. 0.84. All graphs were generated in R ver. 4.0.2 (R Core Team, 2013) with the *ggplot2* package ver. 3.3.3 (Wickham, 2016).

### Supervised Classification Modeling

Supervised classification approaches were used to determine if architectural phenotypes could accurately predict the None/All ratio and root lodging susceptibility. In R ver. 4.0.2 (R Core Team, 2013) the following packages were used for modeling: *tidyverse* ver. 1.3.0 (Wickham et al., 2019), *tidymodels* ver. 0.1.2 (Kuhn and Wickham, 2020), *ranger* ver. 0.12.1 (Wright and Ziegler, 2017), and *kknn* ver. 1.3.1 (Schliep et al., 2016). For each of the models, data was split into a 90% training set and 10% testing set with a 5-fold cross validation strategy.

A multiple regression model was built using architectural phenotypes to predict the None/All ratio and root lodging susceptibility. Four models were tested: 1) the outcome (None/All ratio) and predictors (architectural phenotypes) were assessed within the same year at the individual plant resolution, 2) the outcome (None/All ratio) and predictors (architectural phenotypes) were quantified within the same year at genotypic average resolution, 3) the outcome (None/All ratio) was a multi-year genotypic average and predictors (architectural phenotypes) were individual plant data, and 4) the outcome (lodging susceptibility) was genotypic average and predictors (architectural phenotypes) were individual plant data.

Additionally, genotypes were grouped for categorical classification. For each of the outcomes (the None/All ratio and root lodging susceptibility), data were first scaled and centered before classified into one of three groups - low, average, or high. For each outcome, genotypes were classified as low when they were more than 1 standard deviation below the mean, genotypes were classified as high when they were more than 1 standard deviation above the mean, and genotypes were classified as average when they were within +/- 1 standard deviation of the mean. Four models were tested: 1) the outcome (None/All ratio) and predictors (architectural phenotypes) were assessed within the same year at the individual plant resolution, 2) the outcome (None/All ratio) was a multi-year genotypic average and predictors (architectural phenotypes) were individual plant data, 3) the outcome (lodging susceptibility) was genotypic average and predictors (architectural phenotypes) were individual plant data, and 4) the outcome (lodging susceptibility) was genotypic average and predictors (architectural phenotypes and None/All ratio) were individual plant data. For each categorical classification model, a random forest approach was used with 100 trees. The average accuracy and Receiver Operating Characteristic (ROC) Area Under the Curve (AUC) were extracted for each model. Phenotypes with the highest mean decrease in Gini were considered the most important predictors of the model (Menze et al., 2009).

### Data Availability

All raw data, processed data, and scripts to analyze data are available on GitHub: https://github.com/EESparksLab/Hostetler_et_al_2021

## Results

### The brace root contribution to anchorage varies among genotypes

We have previously shown that brace roots directly contribute to anchorage by establishing a rigid base and that plants with more brace root whorls in the ground have a greater contribution to anchorage (Reneau et al., 2020). However, this report included one genotype with limited phenotypic classification. In this study, we hypothesized that brace roots will have variable contribution to anchorage due to genotype and tested this hypothesis by measuring the None/All ratio (Reneau et al., 2020) for 52 maize inbred lines across two years (Table S1). The None/All ratio quantifies the reduction in anchorage upon removal of brace roots. Thus, a high None/All ratio (close to 1) shows that brace roots have limited impact on anchorage, whereas a low None/All ratio (close to 0) shows that anchorage is dependent on the brace roots in the ground. The None/All ratio data was analyzed by an ANOVA and the effect of genotype (p<0.001), but not year (p>0.05), was significant (Figure 1, Table S3-S4). Thus, confirming our hypothesis that the contribution of brace roots to anchorage varies among genotypes.

**Figure 1.**
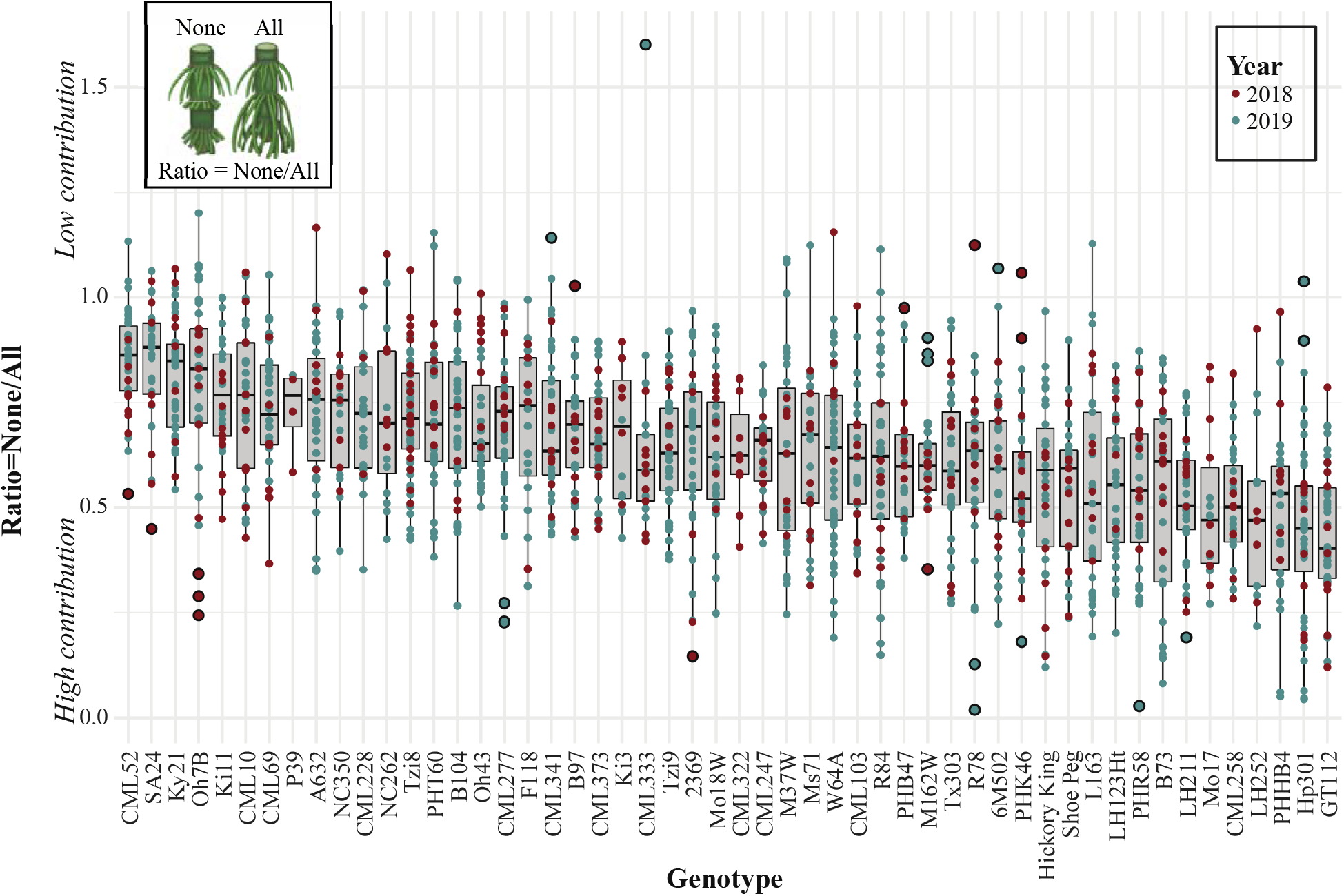
The brace root contribution to anchorage varies depending on genotype. A two-way analysis of variance (ANOVA) showed that the None/All ratio was impacted by genotype (p<0.05) but not year (p>0.05). A higher None/All ratio indicates that anchorage was not influenced by brace roots, whereas a lower None/All ratio indicates that anchorage is dependent on brace roots. Genotypes on the left have an overall higher None/All ratio, whereas genotypes on the right have a lower None/All ratio. The color of dot illustrates the year the data was collected. Outliers are outlined in black.

In support of the hypothesis that the contribution of brace roots to anchorage is genetically controlled, the broad sense heritability (H^2^) estimate was 0.30 (Table S5). However, when considering population structure, the None/All ratio was identified as a homoplasious trait (Figure S4). For example, Hp301 ranked high (51) and SA24 ranked low (2) for the comparative average None/All ratio across years (Figure 1), yet both genotypes are popcorn and cluster phylogenetically (Figure S4). Collectively, these findings establish that the contribution of brace roots to anchorage is dependent on genotype but is not inherited from a common ancestor.

### Plant phenotypes vary among genotypes

We hypothesized that the variation in the brace root contribution to anchorage (None/All ratio) among genotypes is due to underlying variation in brace root phenotypes. To link brace root phenotypes to the brace root contribution to anchorage necessitates a high throughput phenotyping workflow. Current root phenotyping methods are either labor intensive, rely on subjective manual data collection, and/or are not targeted for brace root phenotyping (Trachsel et al., 2011; Das et al., 2015; Zhan et al., 2019; Salungyu et al., 2020, Sanguineti et al., 1998; Liu et al., 2012; Sharma and Carena, 2016). Thus, we developed a workflow that non-destructively captures RGB (Red Green Blue) images using a ground-based robot and extracts brace root phenotypes through a semi-automated root tagging pipeline (Figure S2). In addition to the phenotypes extracted with this pipeline, brace root whorl number in the ground, and plant height were manually quantified for the 52 genotypes to provide an assessment of plant architecture for one year (Figure 2A; Table S6).

**Figure 2.**
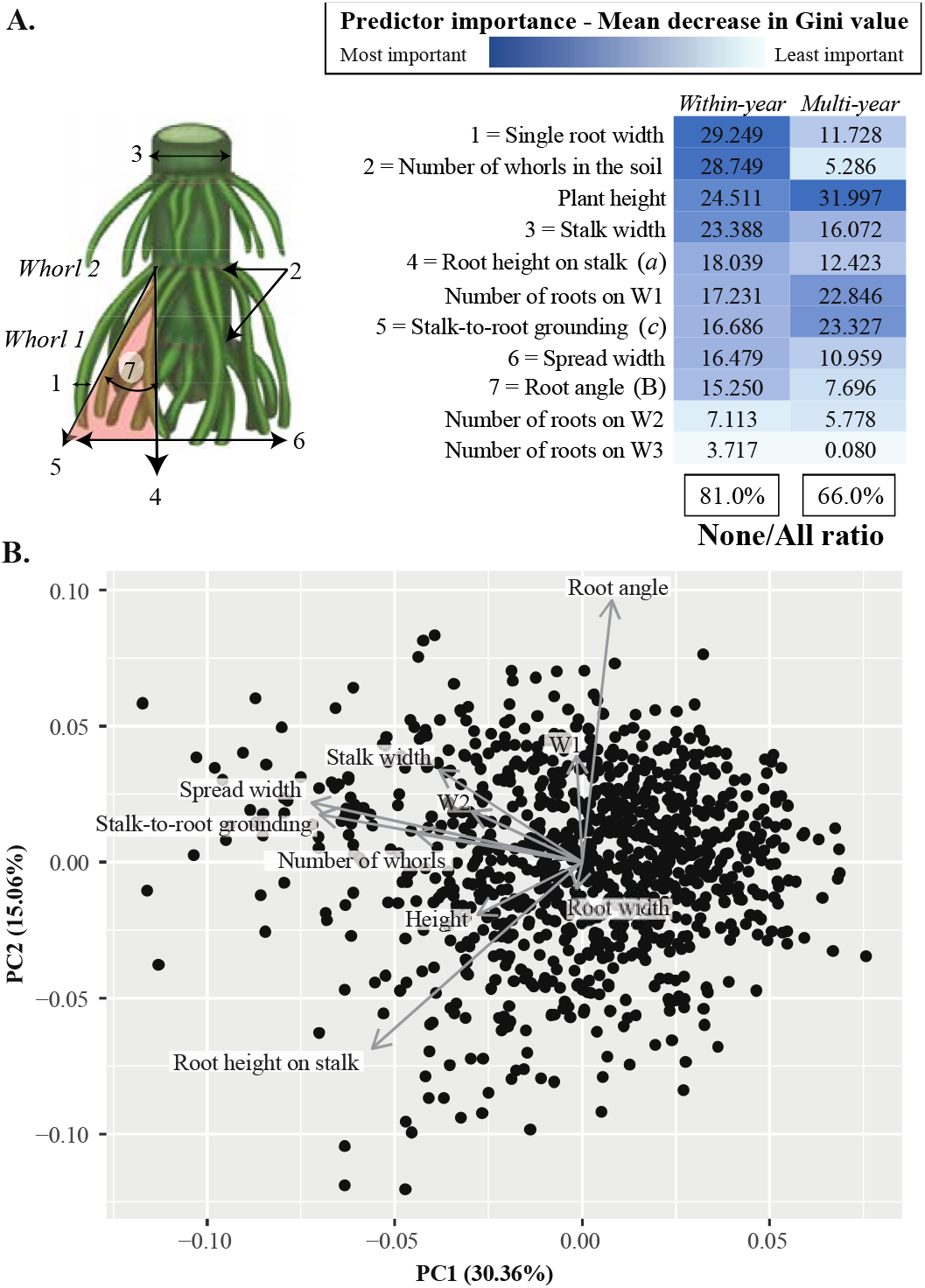
Plant architecture varies among genotype and multiple architectures are important in the contribution of brace roots to anchorage. (A) Brace root phenotypes were quantified with a semi-automated root tagging workflow and phenotypes were only extracted from the brace root whorls that entered the ground. Phenotypes 1, 4, 5, 6, and 7 were extracted for the top-most whorl that entered the ground. The number of roots on each whorl was quantified for all brace root whorls that entered the ground with W1 being the bottom-most whorl (closest to the ground) and W3 being the top-most whorl. The red triangle highlights the right triangle that is used to identify the height of the whorl (*a* leg of the right triangle), the stalk-to-root grounding (*c* leg of the right triangle), and the root angle (B angle of the right triangle). Brace root phenotypes and plant height were used as predictors in random forest models. The predictor importance for each of the random forest models was determined by the mean decrease in Gini value. A darker shade of blue indicates that the corresponding predictor had a higher mean decrease in Gini value and thus was more important in the decision tree, whereas a lighter shade of blue/white indicates that the corresponding predictor had a lower mean decrease in Gini value and thus was less important. Boxed percentages indicate the prediction accuracy for the corresponding model. (B) A principal component analysis (PCA) showed a dense, continuous distribution of plants by phenotype, with the first principal component (PC1) explaining 30.36% of the variation and the second principal component (PC2) explaining 15.06% of the variation. PC1 is primarily driven by spread width and the stalk-to-root grounding, whereas PC2 is primarily driven by root angle.

All plant architectures that were measured varied significantly by genotype (Figures S5, Tables S7-S8). A Principal Component Analysis (PCA) showed a dense, continuous distribution of plants suggesting that a single phenotype was not driving distinct groups or clusters of plants (Figure 2B). The first dimension of the PCA explains 30.40% of the variation and was primarily driven by spread width and the stalk-to-root grounding (Tables S9-S10). The next 6 dimensions explain an additional 64.10% of the variation (>90% total variation explained) and were each driven by a single phenotype (Tables S9-S10). As expected, phenotypes that were highly correlated (Figure S6) also had eigenvectors that loaded in the same plane in the PCA (Figure 2B). For example, the spread width and the stalk-to-root grounding eigenvectors load in the same plane and direction (Figure 2B) and were highly positively correlated (r = 0.99; Figure S6). There is also a negative correlation between the root angle and root height on stalk (r = -0.61; Figure S6) and eigenvectors load in the same plane but opposite directions (Figure 2B). Interestingly, this relationship shows that brace root whorls that attach higher on the stalk have a more acute angle. Consistent with the genetic regulation of these phenotypes, H^2^ for the majority of phenotypes was greater than 0.25 with the highest H^2^ at 0.78 for plant height (Table S5). Thus, these data demonstrate that there is underlying variation in individual plant architectures that may contribute to variation in the brace root contribution to anchorage.

### Multiple phenotypes predict the contribution of brace roots to anchorage

Analysis of the relationship between individual plant architectures and the brace root contribution to anchorage show overall moderate correlations (Table 1). Consistent with previous results (Liu et al., 2012; Sharma and Carena, 2016; Shi et al., 2019; Reneau et al., 2020), the highest correlation was between the number of brace root whorls that entered the ground and the None/All ratio for within-year data (r = -0.41; Table 1). Interestingly, when phenotype data was compared to multi-year None/All ratio data, the highest correlation was with the stalk-to-root grounding (r = -0.36; Table 1). Overall, the correlations between plant architectures and the None/All ratio were higher for within-year comparisons than for between year comparisons. However, even for within-year comparisons, the correlations are moderate suggesting that individual phenotypes cannot explain the contribution of brace roots in anchorage.

**Table 1.** Individual phenotype correlations with the None/All ratio and root lodging susceptibility. A Pearson correlation was run on genotypic averages of plant architectures, the None/All ratio, and root lodging susceptibility. Data represents r correlation values.

|  | Within-year | Multi-year | Lodging<br>susceptibility |
| --- | --- | --- | --- |
|  | None/All ratio | None/All ratio |  |
| Stalk-to-root grounding | -0.37 | -0.361 | -0.193 |
| Number of whorls in ground | -0.41 | -0.355 | -0.433 |
| Plant height | -0.233 | -0.315 | 0.313 |
| Root angle | -0.358 | -0.315 | -0.282 |
| Spread width | -0.318 | -0.310 | -0.174 |
| Stalk width | 0.265 | 0.259 | 0.09 |
| Number of roots on W2 | -0.199 | -0.239 | -0.286 |
| Single root width | -0.211 | -0.178 | -0.064 |
| Number of roots on W1 | 0.106 | 0.107 | 0.091 |
| Number of roots on W3 | 0.081 | 0.057 | -0.1 |
| Height of whorl | -0.017 | -0.036 | 0.125 |
| None/All ratio (Multi-year) | NA | NA | 0.363 |

To further explore the relationship between plant architectures and the contribution of brace roots to anchorage, machine learning models were developed. First, multiple regression models were used to predict the None/All ratio from the plant architectures. For within-year models, both individual plant and genotypic average data predicted the None/All ratio with an R^2^ value of 0.23 (Table 2). However, when a multi-year genotypic average was used for the None/All ratio, the R^2^ value decreases to 0.19 (Table 2). Thus, within-year predictions are stronger than between year predictions.

**Table 2.**
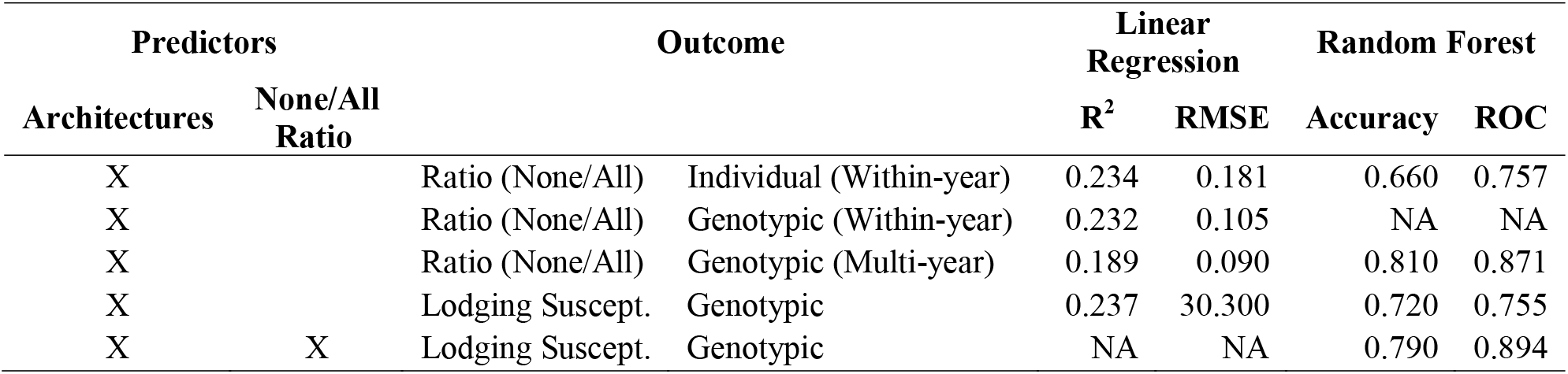
Prediction accuracy of supervised classification models. Supervised classification models (linear regression and random forest models) were used to predict the None/All ratio and lodging susceptibility from plant architectures. The average prediction accuracy and Receiver Operating Characteristic (ROC) Area Under the Curve (AUC) values were extracted from models. Models were run either on individual (within-year) data or genotypic averages for outcomes. The predictors included in each model are indicated with an X.

| Predictors |  | Outcome | Linear<br>Regression |  | Random Forest |  |
| --- | --- | --- | --- | --- | --- | --- |
| Architectures | None/All<br>Ratio |  | R<sup>2</sup> | RMSE | Accuracy | ROC |
| X |  | Ratio (None/All) Individual (Within-year) | 0.234 | 0.181 | 0.660 | 0.757 |
| X |  | Ratio (None/All) Genotypic (Within-year) | 0.232 | 0.105 | NA | NA |
| X |  | Ratio (None/All) Genotypic (Multi-year) | 0.189 | 0.090 | 0.810 | 0.871 |
| X |  | Lodging Suscept. Genotypic | 0.237 | 30.300 | 0.720 | 0.755 |
| X | X | Lodging Suscept. Genotypic | NA | NA | 0.790 | 0.894 |

Next, a categorical classification approach was used to determine if plant architectures can predict the relative classification (low, average, high) of the None/All ratio. Random forest models were generated using the individual plant architectures as predictor variables and the classification of within-year individual plant data or multi-year genotype average data as the outcome. From these models, the None/All ratio category (low, average, or high) was predicted with 66.00% accuracy (ROC AUC=0.76) for within-year data and 81.00% accuracy (ROC AUC=0.87) for multi-year genotype average data (Table 2). The increase in prediction accuracy when considering genotypic averages is likely due to a reduction in noise that is inherent to each measurement. These data further demonstrate that multi-year genotypic averages perform well for the relative classification of the brace root contribution to anchorage as opposed to individual plant data predicting specific outcomes in the multiple regression.

The predictors that are driving the random forest decision trees in each of the outcome models were investigated to understand the plant architectures that are important for classifying the brace root contribution to anchorage. For random forest models, the mean decrease in Gini value can be used to assess the relative importance of each predictor in the decision tree. A higher Gini value indicates a greater influence on the decision tree and thus a more important predictor. For the random forest model that used within-year data, the top predictors included the single root width, the number of brace root whorls in the ground, and plant height (Figure 2A). For the model that used multi-year genotypic averages, the top predictors included plant height, the stalk-to-root grounding, and the number of roots on whorl 1 (Figure 2A). The influence of multiple phenotypes on prediction accuracy is consistent with the moderate univariate correlations we reported (Table 1) and the inconsistent phenotypes identified as important for root lodging in previous studies (Liu et al., 2012; Sharma and Carena, 2016).

### Genotype determines root lodging susceptibility

In 2020, a natural root lodging event occurred at our Newark, DE field site during the late-vegetative and early reproductive growth stages (65 DAP) after brace roots had entered the ground. Tropical Storm Isaias brought heavy rains followed by sustained winds that averaged 23.8 miles per hour (mph) with gusts up to 59 mph (Table S11). Winds were from the West and Northwest, but root lodging was not influenced by field position relative to the wind (Figure S7). This storm provided the unique opportunity to perform a detailed assessment of root lodging within the same germplasm that we previously assessed for the brace root contribution to anchorage and plant architectures.

Within the genotypes, there was a continuous distribution of root lodging susceptibility, with 826 plants remaining upright while 421 plants root-lodged (Figure 3A). A genotypic-level categorization identified 16 root lodging resistant genotypes, with no root lodging in either plot replicate (Figure 3B), and 36 genotypes that had variable root lodging in one or more plots (Figure 3C). As expected, the genotypes that were classified as lodging resistant had more brace root whorls in the ground compared to those that were considered variable (p<0.05, Figure 3D). The broad sense heritability (H^2^) estimate of root lodging susceptibility was 0.58 (Table S5). However, the phylogenetic relationship of genotypes showed that, like the brace root contribution to anchorage, root lodging resistance is independent of population structure (Figure S4). Together, these data are consistent with the genotype-specific, but not subpopulation specific, susceptibility to root lodging.

**Figure 3.**
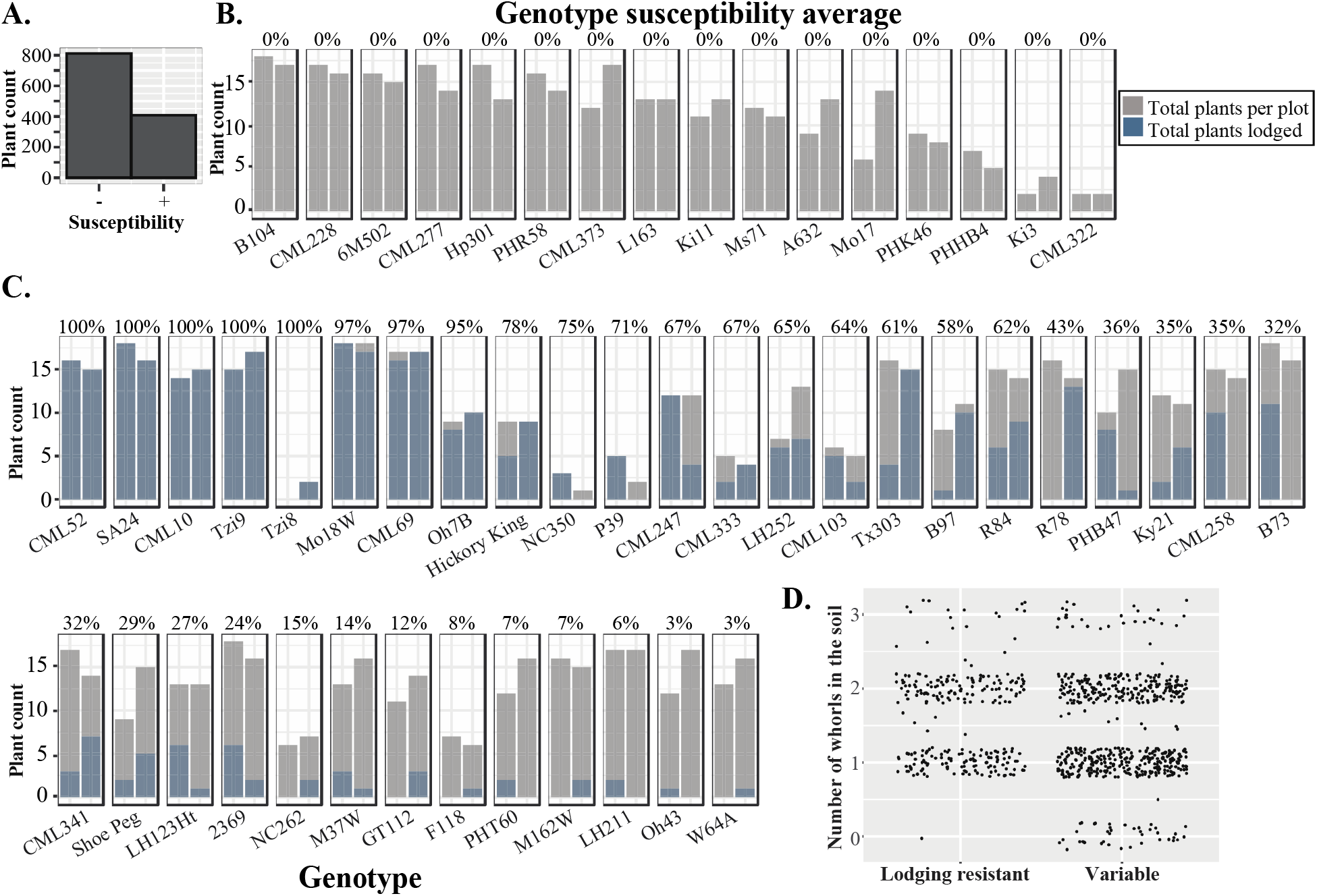
Root lodging varied across 52 maize genotypes. (A) Following Tropical Storm Isaias, 826 plants remained vertical (-) and 421 plants experienced root lodging (+). (B-C) Each bar represents one replicate plot; gray bars indicate the total number of plants per plot; blue bars indicate the total number of plants that lodged per plot; the genotype susceptibility average (shown at the top of both plots) was defined as ‘the total number of plants lodged per genotype/the total number of plants per genotype×100’. (B) There were 16 genotypes that did not root lodge in either plot replicate. (C) There were 36 genotypes that showed variable degrees of root lodging in both plot replicates. (D) The genotypes that were classified as lodging resistant (B) had more brace root whorls in the ground compared to those that were variable (C).

### The contribution of brace roots to anchorage is a good proxy for root lodging susceptibility

To determine if the None/All ratio is an appropriate proxy for identifying lodging resistant genotypes, a pairwise Pearson correlation was calculated. The correlation between the multi-year genotypic averages of the None/All ratio and root lodging susceptibility was surprisingly high (r = 0.36; Table 1) given that the data were from different years and different growth stages. These data provide additional support for the importance of brace roots for anchorage, with the positive correlation showing that the greater the brace root contribution to anchorage (i.e., a lower ratio), the less likely the plant is to root lodge.

To determine if the same phenotypes that are important for the None/All ratio are also important in lodging susceptibility, pairwise Pearson correlations were run (Table 1). Like the correlations between phenotypes and the None/All ratio, there were overall moderate correlations between individual phenotypes and root lodging susceptibility (Table 1). The two highest correlations were with the number of whorls in the ground (r = -0.43) and plant height (r = 0.31). Interestingly, these same phenotypes had high correlations with the None/All ratio between and within-years.

When considering a multiple regression model using individual plant architectures to predict genotypic averages of root lodging susceptibility, the R^2^ value is 0.24 (Table 2). This is the same predictive power as when using the plant architectures to predict the None/All ratio. Thus, these data show that end-of-season plant architectural phenotyping data can be used to predict mid-season root lodging with surprising accuracy.

When the categorical classification approach was used, the relative genotype classification (low, average, high) for root lodging susceptibility was classified with 72.00% accuracy (ROC AUC=0.76; Table 2). With the inclusion of the None/All ratio, the prediction accuracy of root lodging susceptibility increased slightly 78.00% (Table 2). Similar to the models that predicted the None/All ratio (Figure 2A), the top predictor of lodging susceptibility was plant height. However, unlike the None/All ratio, the rest of the plant architectures are approximately equal in their contribution to the decision tree (Figure 4). Overall, this data confirms the utility of multi-year data to provide insight into the plant architectures that promote anchorage and limit root lodging.

**Figure 4.**
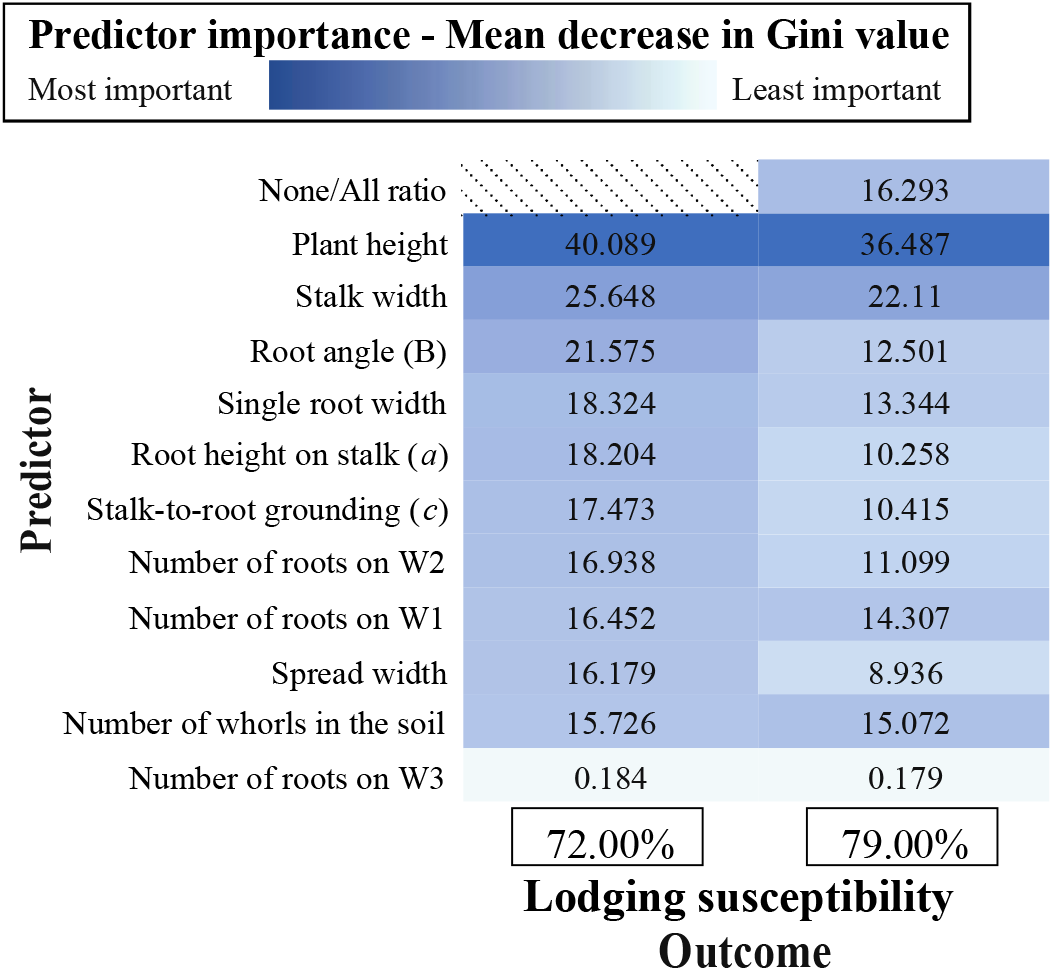
Multiple plant architectures predict root lodging susceptibility. (A) The predictor importance for each of the random forest models was determined by the mean decrease in Gini value. A darker shade of blue indicates that the corresponding predictor had a higher mean decrease in Gini value and thus was more important in the decision tree, whereas a lighter shade of blue/white indicates that the corresponding predictor had a lower mean decrease in Gini value and thus was less important. Boxed percentages indicate the prediction accuracy for the corresponding model. Plant architecture predicted root lodging susceptibility with 72.00% accuracy, whereas the inclusion of the None/All ratio increased prediction accuracy by 7.00%. The top predictor for both models is plant height.

### Plant height is positively correlated with brace root phenotypes but has opposing effects on root lodging susceptibility

Plant height has been historically associated with lodging, as evidenced by the development of dwarf varieties of grain crops during the Green Revolution, which reduced the impact of lodging. The moderate correlations between plant height and lodging susceptibility (r = 0.31) in this study suggest that plant height alone cannot not explain root lodging in maize (Table 1). Interestingly, there were positive correlations between brace root phenotypes and plant height (Figure S6), but they have opposing effects on lodging susceptibility (Table 1). In other words, taller plants are more likely to root lodge, but taller plants also have larger brace root traits, which limits root lodging. When genotypic averages were used, plant height was universally the most important predictor (Figure 2A, Figure 4) in machine learning models. Together, these data show that the contribution of multiple phenotypes is important for limiting root lodging and highlights the importance of uncoupling plant height and root system traits for future plant improvement.

## Discussion

This study was motivated by a limited understanding of how plant architectures contribute to plant anchorage and root lodging resistance. Using a multi-year data acquisition strategy, we assessed 1) the brace root contribution to root anchorage, 2) plant architectural phenotypes, and 3) root lodging susceptibility in a population of 52 maize inbred lines. Surprisingly, we found that multi-year genotypic-level data were highly successful in predictive, categorical modeling.

In linking plant architectures to anchorage and root lodging resistance, we demonstrate that plant height is important for both outcomes. Generally, genotypes classified as root lodging resistant were shorter and with a greater brace root contribution to anchorage than genotypes that were classified as root lodging susceptible. This is consistent with the prevailing idea from the Green Revolution that short plants are more lodging-resistant. However, the correlation was moderate (r = 0.31; Table 1), which indicates that plant height is not the only factor influencing root lodging. Indeed, this is consistent with the variable reports on plant height and lodging in maize (Brune et al., 2018; Sharma and Carena, 2016). In addition to plant height, we show brace root phenotypes that collectively establish a wider base are associated with an increased contribution to anchorage and root lodging resistance. Interestingly, these same phenotypes are associated with increased plant height. Thus, a genetic decoupling of plant architectures is necessary to fully optimize plants for root lodging-resistance.

Plant anchorage is an outcome of both above- and below-ground root systems, and their interaction with the ground. Despite this complex relationship, we find that genotypes with a higher brace root contribution to anchorage are more likely to resist lodging. This result demonstrates that either brace roots are the primary contributor to anchorage, or the functional and phenotypic assessment of the above ground root system is reflective of the function and phenotype of the below ground.

## Abbreviations

AUC: Area Under the Curve
BRobot: Brace Root phenotyping robot
DARLING: Device for Assessing Resistance to Lodging IN Grains
ROC: Receiver Operating Characteristic

## Acknowledgements

We thank members of the Sparks lab, Sarah Blizard and Noah Ouslander, for assistance in mechanical testing. We thank William Cantera (University of Delaware) for assistance in developing, Dr. Jason Wallace (University of Georgia) for assistance in updating, and the Vertically Integrated Projects course for beta-testing and debugging of the root tagging pipeline. We thank Dr. Rajdeep Khangura (Purdue University) and Dr. Brandon Sinn (Otterbein University) for input in data analysis. We thank Teclemariam Weldekidan (University of Delaware), Dr. Colleen Drapek (Sainsbury Lab Cambridge), Dr. Anjali Iyer-Pascuzzi (Purdue University), Dr. Brian Dilkes (Purdue University) and Dr. Kevin Lehner (Colorado State University) for providing manuscript feedback. This work was supported by a Delaware Center for Advanced Technology grant and a University of Delaware Research Foundation (UDRF) award to EES.

## Author Contributions

JWR, LE, AS, and EES collected data. ANH and EES analyzed data. AS and HGT developed the semi-automated root phenotyping workflow. DDS provided DARLING devices and trained operators in their use. All authors contributed to the writing and/or editing of the manuscript.

## Supplementary Data

**Figure S1. A ground-based brace root phenotyping robot (BRobot) was used for image capture**. BRobot is a modified Superdroid LT2 Tracked ATR Robot Platform with a custom controller. The red arrow highlights the side-mounted FLIR 3.2 MP Color Blackfly camera that captures images.

**Figure S2. A semi-automated root tagging workflow was developed to optimize image processing**. After RGB images are acquired, a python script (*main*.*py*) launches the root tagging graphical user interface (GUI). A tag of known size (0.5 inches) was placed on the stem prior to image acquisition for scale. First, the tag is selected for a pixel (px) scale, then each additional screen will prompt the user to click and/or record/identify specific regions in the image. The red dots shown on the image highlight the following phenotypes that have been tagged: 1) the number of pixels within a 0.5-inch region, 2) the stalk width, and 3) the single root width. The purple dots shown on the image highlight the right triangle that is used to identify the following phenotypes: 1) the height of the whorl, 2) the stalk-to-root grounding, and 3) the root angle. The number of roots within each whorl was counted and typed into the white boxes (shown on the left). This records the total number of roots per whorl and the number of whorls in the ground. After recording or identifying phenotypes, the user will hit “enter.” After all phenotypes have been recorded, the user will hit “Next” (red bar on the right of the screen) to begin the next image. After completing all images, a python script (*process_rootpixel_data*.*py*) is used to convert pixel data to scaled phenotype data. All data is exported to a .csv file for processing and analysis.

**Figure S3. Phenotypes extracted from left and right images were highly correlated**. Left and right images (labeled A and B) were acquired for each plant and phenotypes were extracted from images with the root tagging GUI. A Pearson correlation analysis was run to determine the precision of our root tagging GUI. Cells highlighted in yellow indicate the same phenotype from both sides of the plant.

**Figure S4. Root lodging is not monophyletic**. A species tree was generated for 41 of the 52 maize inbred genotypes included in this study. Genotype names are color coded according to their assigned subpopulation information [Flint-Garcia et al (2005) and Liu et al (2003)]. If all genotypes within a clade are from the same subpopulation, branches are solid and colored with the respective subpopulation. Branches are dotted if the clade includes genotypes from more than one subpopulation. Genotypes were identified as lodging resistant or variable. The assigned lodging classification is illustrated to the right of the genotype with a circle or square. Clades where all genotypes within the clade are a part of a single lodging classification are highlighted with the corresponding shape at the node. Numbers to the right of the lodging category indicate the rank order of the None/All ratio from Figure 1, where 1 indicates genotypes that have a high None/All ratio and 52 indicates genotypes that have a low None/All ratio.

**Figure S5. Plant phenotypes vary among genotypes**. The following phenotypes vary among genotypes: (A) The number of roots on whorl 1 (the whorl closest to the ground, bottom whorl), (B) the number of roots on whorl 2 (middle whorl), (C) the number of roots on whorl 3 (top whorl), (D) the single root width, (E) the stalk width, (F) the root height on stalk, (G) the stalk-to-root grounding, (H) the root angle, (I) the spread width, (J) the number of whorls in the ground, and (K) plant height. (A-K) Genotypes are ordered by rank (high to low) for the None/All ratio according to Figure 1. The color of each dot illustrates the replicate plot where the phenotype data is from. Outliers are outlined in black.

**Figure S6. Brace root phenotypes are correlated with each other**. A Pearson correlation analysis shows which phenotypes are highly correlated.

**Figure S7. Root lodging was not related to field position**. Winds from the West and North-West induced root lodging. Plots were colored according to the susceptibility of root lodging within each plot. Plots with 100% root lodging were highlighted with white, whereas plots that had 0% root lodging were highlighted with dark red. Plots with a grid overlaid indicate plots that did not germinate.

**Table S1. List of 52 maize inbred genotypes with assigned subpopulation information**. The subpopulation information for each of the 52 maize inbred genotypes included in this study was identified in Flint-Garcia et al (2005) and Liu et al (2003). Genotypes with missing information are denoted with a (*).

**Table S2. Temperature, rainfall, and wind speed summaries for the 2018, 2019, and 2020 field seasons**. For the duration of this study, the average temperature (degrees F), total rainfall (in), and average wind speeds (mph) were included for each of the months in the 2018, 2019, and 2020 field seasons. Additional weather data can be found at the Delaware Environmental Observing System (http://www.deos.udel.edu) and selecting the Newark, DE-Ag Farm station.

**Table S3. Analysis of variance (ANOVA) results for the None/All ratio**. A two-way ANOVA was run for the None/All ratio with genotype and year included as the independent variables. The None/All ratio varied among genotypes (p<0.001) but not years (p>0.05).

**Table S4**. Pairwise comparisons for 2018 and 2019 plant biomechanics. A post-hoc Tukey’s Honest Significant Difference (HSD) test was used to determine genotypes that were significantly different for the None/All ratio. Genotypes that share a letter indicate genotypes that are not significantly different, whereas genotypes that do not share a letter are significantly different. The p-value was assessed at <0.05.

**Table S5. Heritability estimates for the None/All ratio, plant phenotype data, and root lodging data**. Broad-sense heritability (H2) was estimated for the None/All ratio, plant phenotype data, and root lodging data.

**Table S6. Descriptions of the data collected within this study**. The trait name, unit of measurement, and description were included for the data quantified within this study.

**Table S7. Analysis of variance (ANOVA) results for plant phenotype data**. A one-way ANOVA was run for each of the plant phenotypes included in this study with genotype as the independent variable and the corresponding plant phenotype as the dependent variable. With the exception of stalk width, all of the phenotypes were transformed with a Tukey ladder of powers transformation prior to running the ANOVA to meet the normality assumption. p<0.001 (***); p<0.01 (**); p<0.05 (*)

**Table S8. Pairwise comparisons for each of the 2019 phenotypes**. A post-hoc Tukey’s Honest Significant Difference (HSD) test was used to determine genotypes that were significantly different for each of the phenotypes measured. Genotypes that share a letter for a single phenotype indicate genotypes that are not significantly different, whereas genotypes that do not share a letter are significantly different. The p-value was assessed at <0.05.

**Table S9. Summary statistics for each of the 10 dimensions from a principal component analysis (PCA)**. A standard deviation and proportion of variance were extracted for each of the 10 dimensions of the PCA. Within the first two dimensions 45.4% of the variance was explained and within seven dimensions more than 90% of the variance was explained.

**Table S10. Variance contributions for each of the 10 dimensions from a principal component analysis (PCA)**. The contribution of each phenotype for dimensions 1-10 was extracted from the PCA.

**Table S11. Environmental Conditions Summary for Tropical Storm Isaias**. The following 24-hour weather summary for August 04, 2020, was collected from the DEOS for the Newark, DE Agriculture Farm station. The Newark, DE Agricultural Farm Station is located at 39° 40’ N and 75° 45’ W at an elevation of 106 ft. Additional data can be viewed at: http://www.deos.udel.edu and selecting the Newark, DE-Ag Farm station.

## Literature Cited

Berry, P. M., M. Sterling, J. H. Spink, C. J. Baker, R. Sylvester-Bradley, S. J. Mooney, A. R. Tams, and A. R. Ennos. 2004. Understanding and reducing lodging in cereals. Advances in Agronomy, 217–271. Elsevier.

Blizard, S., and E. E. Sparks. 2020. Maize Nodal Roots. Annual Plant Reviews online, 281–304. American Cancer Society.

Brune, P. F., A. Baumgarten, S. J. McKay, F. Technow, and J. J. Podhiny. 2018. A biomechanical model for maize root lodging. Plant and Soil 422: 397–408.

Carter, P. R., and K. D. Hudelson. 1988. Influence of simulated wind lodging on corn growth and grain yield. Journal of Production Agriculture 1: 295–299.

Chifman, J., and L. Kubatko. 2015. Identifiability of the unrooted species tree topology under the coalescent model with time-reversible substitution processes, site-specific rate variation, and invariable sites. Journal of Theoretical Biology 374: 35–47.

Chifman, J., and L. Kubatko. 2014. Quartet Inference from SNP Data Under the Coalescent Model. Bioinformatics 30: 3317–3324.

Cook, D. D., W. de la Chapelle, T.-C. Lin, S. Y. Lee, W. Sun, and D. J. Robertson. 2019. DARLING: a device for assessing resistance to lodging in grain crops. Plant methods 15: 102.

Das, A., H. Schneider, J. Burridge, A. K. M. Ascanio, T. Wojciechowski, C. N. Topp, J. P. Lynch, et al. 2015. Digital imaging of root traits (DIRT): a high-throughput computing and collaboration platform for field-based root phenomics. Plant Methods 11: 51.

Dhugga, K. S. 2007. Maize Biomass Yield and Composition for Biofuels. Crop Science 47: 2211–2227.

Erndwein, L., D. D. Cook, D. J. Robertson, and E. E. Sparks. 2020. Field-based mechanical phenotyping of cereal crops to assess lodging resistance. Applications in Plant Sciences 8: e11382.

Farkhari, M., A. Krivanek, Y. Xu, T. Rong, M. R. Naghavi, B. Y. Samadi, and Y. Lu. 2013. Root-lodging resistance in maize as an example for high-throughput genetic mapping via single nucleotide polymorphism-based selective genotyping. Plant Breeding 132: 90–98.

FlintLGarcia, S. A., C. Jampatong, L. L. Darrah, and M. D. McMullen. 2003. Quantitative Trait Locus Analysis of Stalk Strength in Four Maize Populations. Crop Science 43: 13–22.

Holland, J. B., W. E. Nyquist, and C.T. CervantesLMartínez. 2002. Estimating and Interpreting Heritability for Plant Breeding: An Update. Plant Breeding Reviews, 9–112. John Wiley & Sons, Ltd.

Hoppe, D. C., M. E. McCully, and C. L. Wenzel. 1986. The nodal roots of Zea: their development in relation to structural features of the stem. Canadian Journal of Botany 64: 2524–2537.

Hostetler, A. N., R. S. Khangura, B. P. Dilkes, and E. E. Sparks. 2021. Bracing for sustainable agriculture: the development and function of brace roots in members of Poaceae. Current Opinion in Plant Biology 59: 101985.

Kassambara, A., and F. Mundt. 2020. factoextra: Extract and Visualize the Results of Multivariate Data Analyses.

Kuhn, M., and H. Wickham. 2020. Tidymodels: a collection of packages for modeling and machine learning using tidyverse principles.

Liu, S., F. Song, F. Liu, X. Zhu, and H. Xu. 2012. Effect of planting density on root lodging resistance and its relationship to nodal root growth characteristics in maize (Zea mays L.). Journal of Agricultural Science 4: 182.

Mangiafico, S. 2021. rcompanion: Functions to Support Extension Education Program Evaluation.

Menze, B. H., B. M. Kelm, R. Masuch, U. Himmelreich, P. Bachert, W. Petrich, and F. A. Hamprecht. 2009. A comparison of random forest and its Gini importance with standard chemometric methods for the feature selection and classification of spectral data. BMC Bioinformatics 10: 213.

R Core Team. 2013. R: A language and enviornment for satistical computing. R Foundation for Statistical Computing, Vienna, Austria.

Rajkumara, S. 2008. Lodging in cereals – a review. Agricultural Reviews 29: 55–60.

Reneau, J. W., R. S. Khangura, A. Stager, L. Erndwein, T. Weldekidan, D. D. Cook, B. P. Dilkes, and E. E. Sparks. 2020. Maize brace roots provide stalk anchorage. Plant Direct 4: e00284.

Robertson, D. J., M. Julias, S. Y. Lee, and D. D. Cook. 2017. Maize Stalk Lodging: Morphological Determinants of Stalk Strength. Crop Science 57: 926–934.

Salungyu, J., S. Thaitad, A. Bucksch, J. Kengkanna, and P. J. Saengwilai. 2020. From lab to field: Open tools facilitating the translation of maize root traits. Field Crops Research 255: 107872.

Sanguineti, M. C., M. Giuliani, G. Govi, R. Tuberosa, and P. Landi. 1998. Root and shoot traits of maize inbred lines grown in the field and in hydroponic culture and their relationships with root lodging. Maydica 43: 211–216.

Schliep, K., K. Hechenbichler, and A. Lizee. 2016. kknn: Weighted k-Nearest Neighbors.

Sharma, S., and M. J. Carena. 2016. BRACE: A Method for High Throughput Maize Phenotyping of Root Traits for Short-Season Drought Tolerance. Crop Science 56: 2996–3004.

Shi, J., B. J. Drummond, J. E. Habben, N. Brugire, B. P. Weers, S. M. Hakimi, H. R. Lafitte, et al. 2019. Ectopic expression of ARGOS8 reveals a role for ethylene in root-lodging resistance in maize. The Plant Journal 97: 378–390.

Stamp, P., and C. Kiel. 1992. Root morphology of maize and its relationship to root lodging. Journal of Agronomy and Crop Science 168: 113–118.

Thompson, D. L. 1972. Recurrent Selection for Lodging Susceptibility and Resistance in Corn1. Crop Science 12: cropsci1972.0011183X001200050023x.

Tirado, S. B., C. N. Hirsch, and N. M. Springer. 2020. Utilizing Temporal Measurements from UAVs to Assess Root Lodging in Maize and its Impact on Productivity. Plant Biology.

Trachsel, S., S. M. Kaeppler, K. M. Brown, and J. P. Lynch. 2011. Shovelomics: high throughput phenotyping of maize (Zea mays L.) root architecture in the field. Plant and Soil 341: 75–87.

Wascher, M., and L. Kubatko. 2021. Consistency of SVDQuartets and Maximum Likelihood for Coalescent-Based Species Tree Estimation. Systematic Biology 70: 33–48.

Wei, T., and V. Simko. 2017. R package ‘corrplot’: Visualization of a Correlation Matrix.

Wickham, H. 2016. ggplot2: Elegant Graphics for Data Analysis. R Foundation for Statistical Computing, Verlag, New York.

Wickham, H., M. Averick, J. Bryan, W. Chang, L. McGowan, R. François, G. Grolemund, et al. 2019. Welcome to the Tidyverse. Journal of Open Source Software 4: 1686.

Wright, M. N., and A. Ziegler. 2017. ranger□: A Fast Implementation of Random Forests for High Dimensional Data in C++ and R. Journal of Statistical Software 77.

Zhan, A., J. Liu, S. Yue, X. Chen, S. Li, and A. Bucksch. 2019. Architectural and anatomical responses of maize roots to agronomic practices in a semi-arid environment. Journal of Plant Nutrition and Soil Science 182: 751–762.

Zhang, Q., F. A. Pettolino, K. S. Dhugga, J. A. Rafalski, S. Tingey, J. Taylor, N. J. Shirley, et al. 2011. Cell Wall Modifications in Maize Pulvini in Response to Gravitational Stress. Plant Physiology 156: 2155–2171.

