## Supplementary Figures for "Multiple brace root phenotypes promote anchorage and limit root lodging in maize"

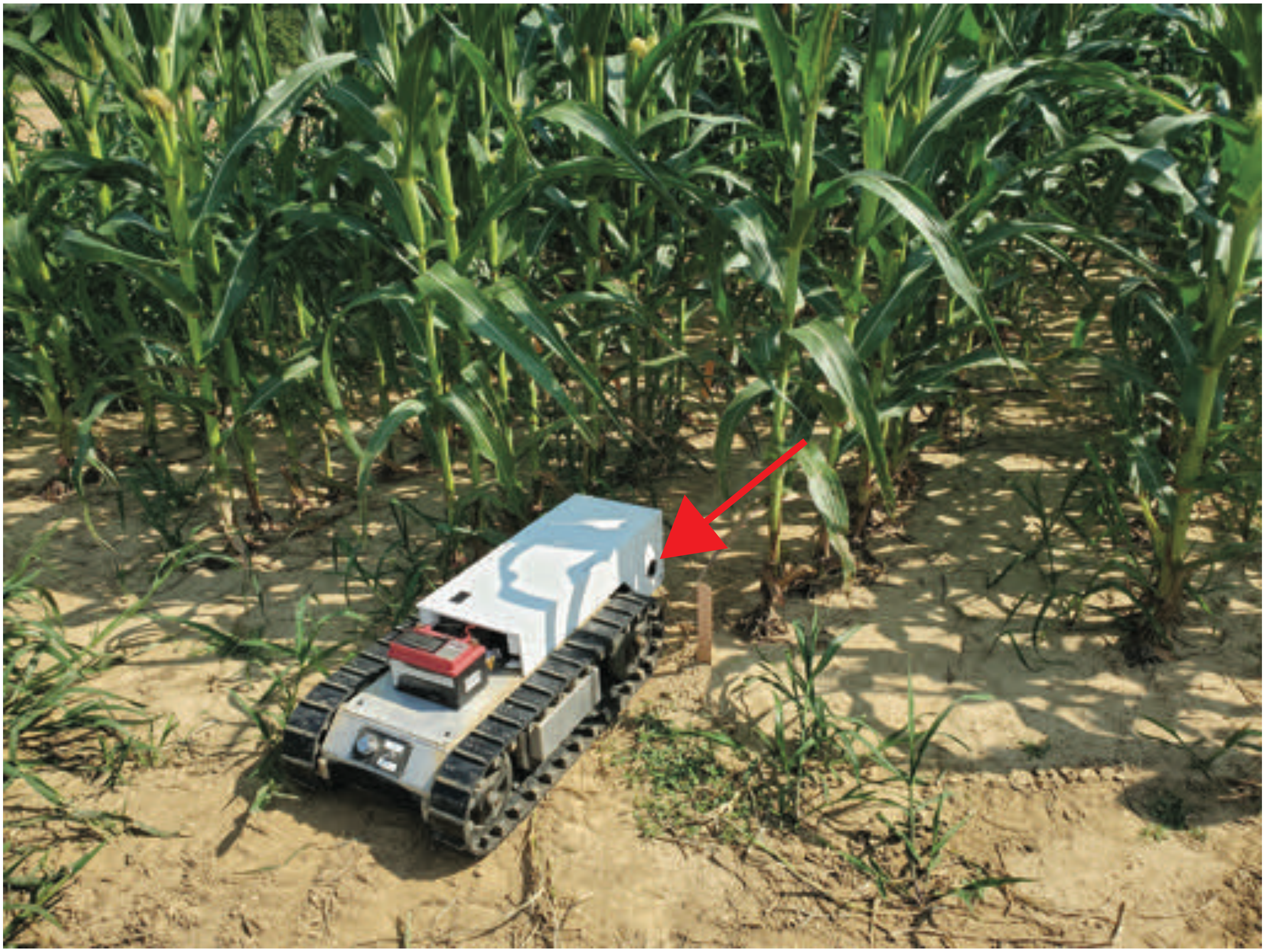

**Figure S1. A ground-based brace root phenotyping robot (BRobot) was used for image capture.** BRobot is a modified Superdroid LT2 Tracked ATR Robot Platform with a custom controller. The red arrow highlights the side-mounted FLIR 3.2 MP Color Blackfly camera that captures images.

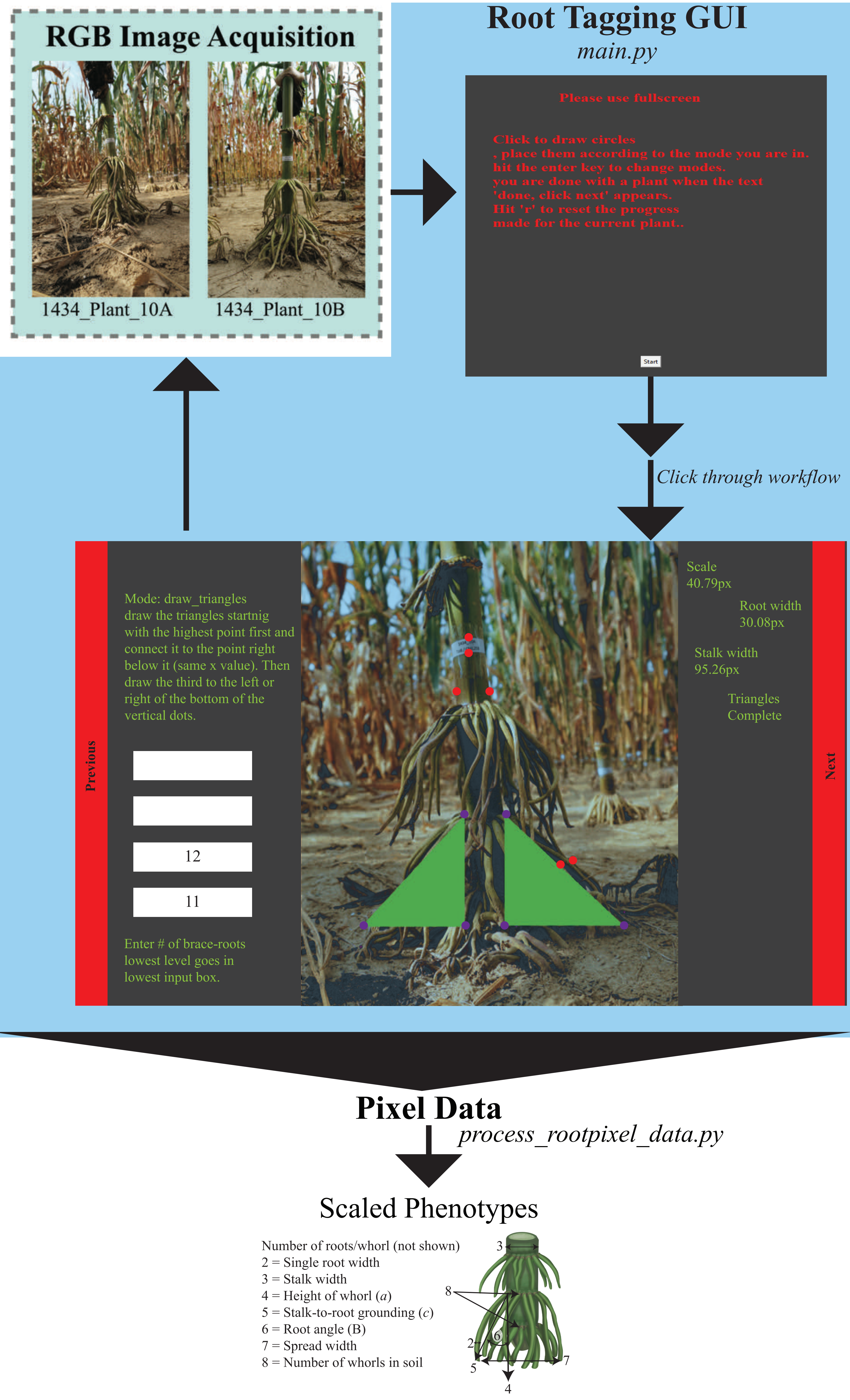
RGB Image Acquisition
1434_Plant_10A
1434_Plant_10B
Root Tagging GUI
main.py
Please use fullscreen
Click to draw circles
, place them according to the mode you are in.
hit the enter key to change modes.
you are done with a plant when the text
'done, click next' appears.
Hit 'r' to reset the progress
made for the current plant..
Start
Click through workflow
Previous
Mode: draw_triangles
draw the triangles startnig
with the highest point first and
connect it to the point right
below it (same x value). Then
draw the third to the left or
right of the bottom of the
vertical dots.
12
11
Enter # of brace-roots
lowest level goes in
lowest input box.
Scale
40.79px
Root width
30.08px
Stalk width
95.26px
Triangles
Complete
Next
Pixel Data
process_rootpixel_data.py
Scaled Phenotypes
Number of roots/whorl (not shown)
2 = Single root width
3 = Stalk width
4 = Height of whorl (a)
5 = Stalk-to-root grounding (c)
6 = Root angle (B)
7 = Spread width
8 = Number of whorls in soil
3
8
2
6
5
7
4

**Figure S2. A semi-automated root tagging workflow was developed to optimize image processing.** After RGB images are acquired, a python script (*main.py*) launches the root tagging graphical user interface (GUI). A tag of known size (0.5 inches) was placed on the stem prior to image acquisition for scale. First, the tag is selected for a pixel (px) scale, then each additional screen will prompt the user to click and/or record/identify specific regions in the image. The red dots shown on the image highlight the following phenotypes that have been tagged: 1) the number of pixels within a 0.5-inch region, 2) the stalk width, and 3) the single root width. The purple dots shown on the image highlight the right triangle that is used to identify the following phenotypes: 1) the height of the whorl, 2) the stalk-to-root grounding, and 3) the root angle. The number of roots within each whorl was counted and typed into the white boxes (shown on the left). This records the total number of roots per whorl and the number of whorls in the ground. After recording or identifying phenotypes, the user will hit "enter." After all phenotypes have been recorded, the user will hit "Next" (red bar on the right of the screen) to begin the next image. After completing all images, a python script (*process_rootpixel_data.py*) is used to convert pixel data to scaled phenotype data. All data is exported to a .csv file for processing and analysis.

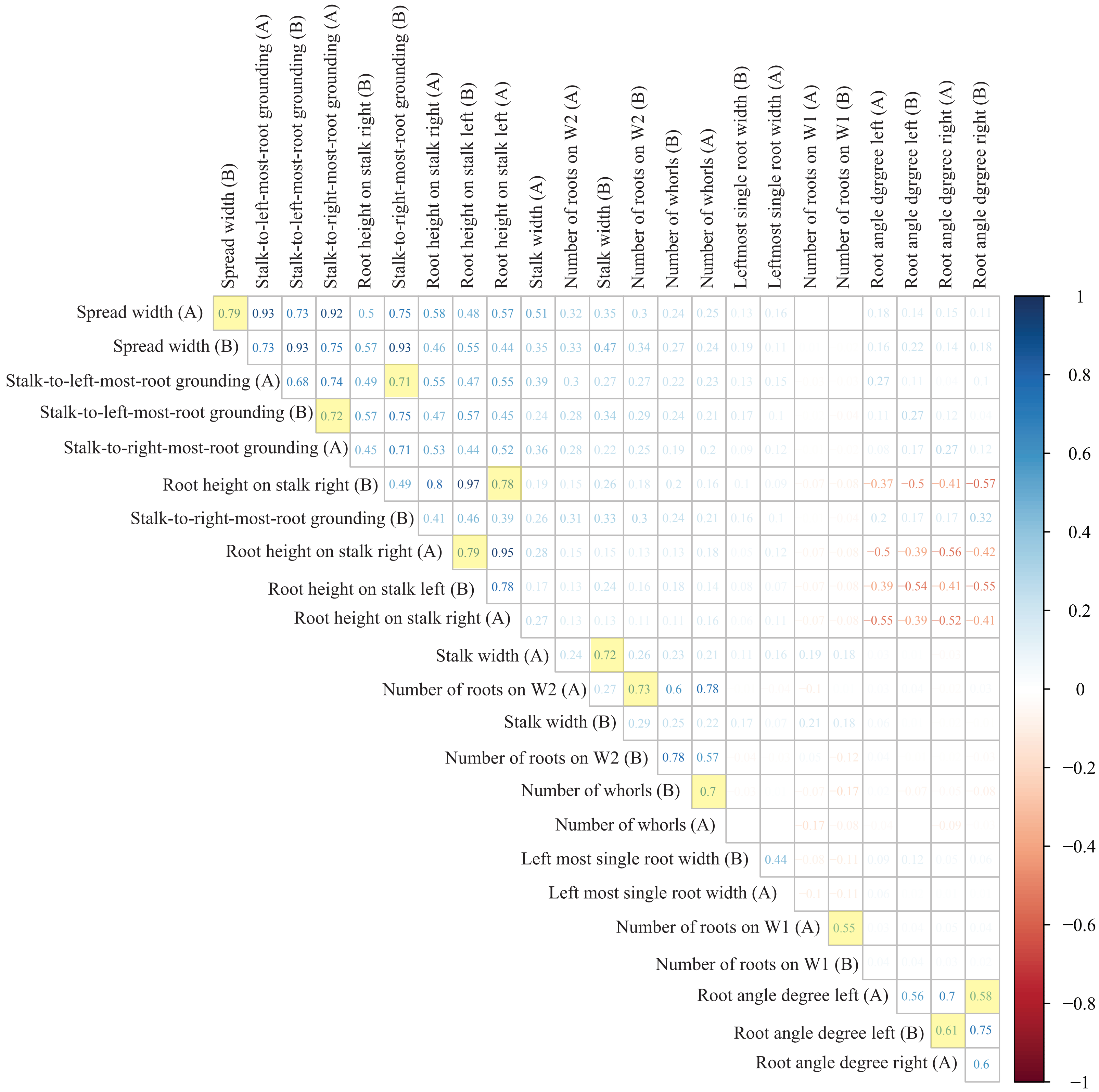
Spread width (B)
Stalk-to-left-most-root grounding (A)
Stalk-to-left-most-root grounding (B)
Stalk-to-right-most-root grounding (A)
Root height on stalk right (B)
Stalk-to-right-most-root grounding (B)
Root height on stalk right (A)
Root height on stalk left (B)
Root height on stalk left (A)
Stalk width (A)
Number of roots on W2 (A)
Stalk width (B)
Number of roots on W2 (B)
Number of whorls (B)
Number of whorls (A)
Leftmost single root width (B)
Leftmost single root width (A)
Number of roots on W1 (A)
Number of roots on W1 (B)
Root angle dgrgree left (A)
Root angle dgrgree left (B)
Root angle dgrgree right (A)
Root angle dgrgree right (B)
Spread width (A)
Spread width (B)
Stalk-to-left-most-root grounding (A)
Stalk-to-left-most-root grounding (B)
Stalk-to-right-most-root grounding (A)
Root height on stalk right (B)
Stalk-to-right-most-root grounding (B)
Root height on stalk right (A)
Root height on stalk left (B)
Root height on stalk right (A)
Stalk width (A)
Number of roots on W2 (A)
Stalk width (B)
Number of roots on W2 (B)
Number of whorls (B)
Number of whorls (A)
Left most single root width (B)
Left most single root width (A)
Number of roots on W1 (A)
Number of roots on W1 (B)
Root angle degree left (A)
Root angle degree left (B)
Root angle degree right (A)
1
0.8
0.6
0.4
0.2
0
−0.2
−0.4
−0.6
−0.8
−1

**Figure S3. Phenotypes extracted from left and right images were highly correlated.** Left and right images (labeled A and B) were acquired for each plant and phenotypes were extracted from images with the root tagging GUI. A Pearson correlation analysis was run to determine the precision of our root tagging GUI. Cells highlighted in yellow indicate the same phenotype from both sides of the plant.

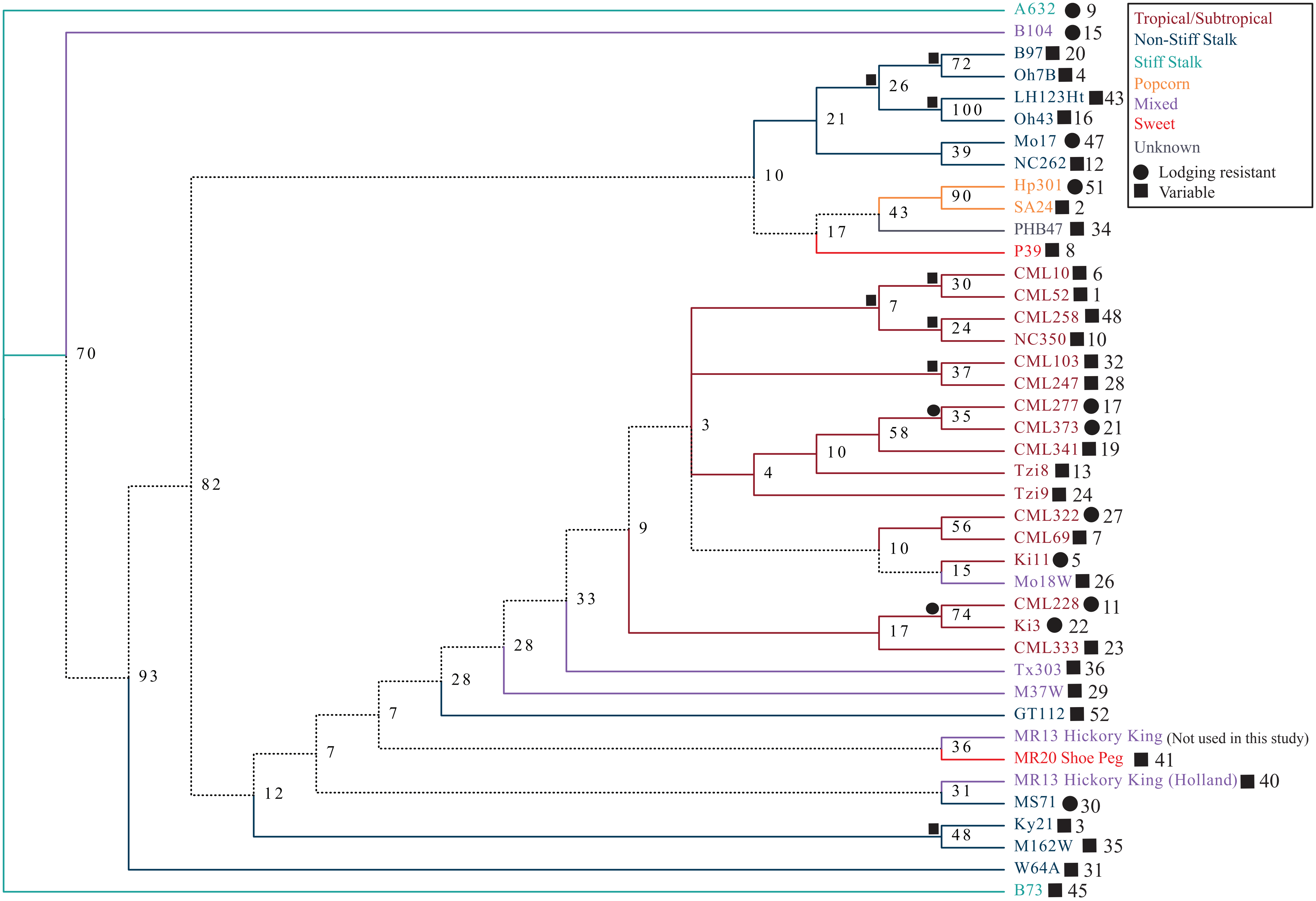


**Figure S4. Root lodging is not monophyletic.** A species tree was generated for 41 of the 52 maize inbred genotypes included in this study. Genotype names are color coded according to their assigned subpopulation information [Flint-Garcia et al (2005) and Liu et al (2003)]. If all genotypes within a clade are from the same subpopulation, branches are solid and colored with the respective subpopulation. Branches are dotted if the clade includes genotypes from more than one subpopulation. Genotypes were identified as lodging resistant or variable.
The assigned lodging classification is illustrated to the right of the genotype with a circle or square. Clades where all genotypes within the clade are a part of a single lodging classification are highlighted with the corresponding shape at the node. Numbers to the right of lodging category indicate rank order of the None/All ratio from Figure 1, where 1 indicates genotypes that have a high None/All ratio and 52 indicates genotypes that have a low None/All ratio.

A.

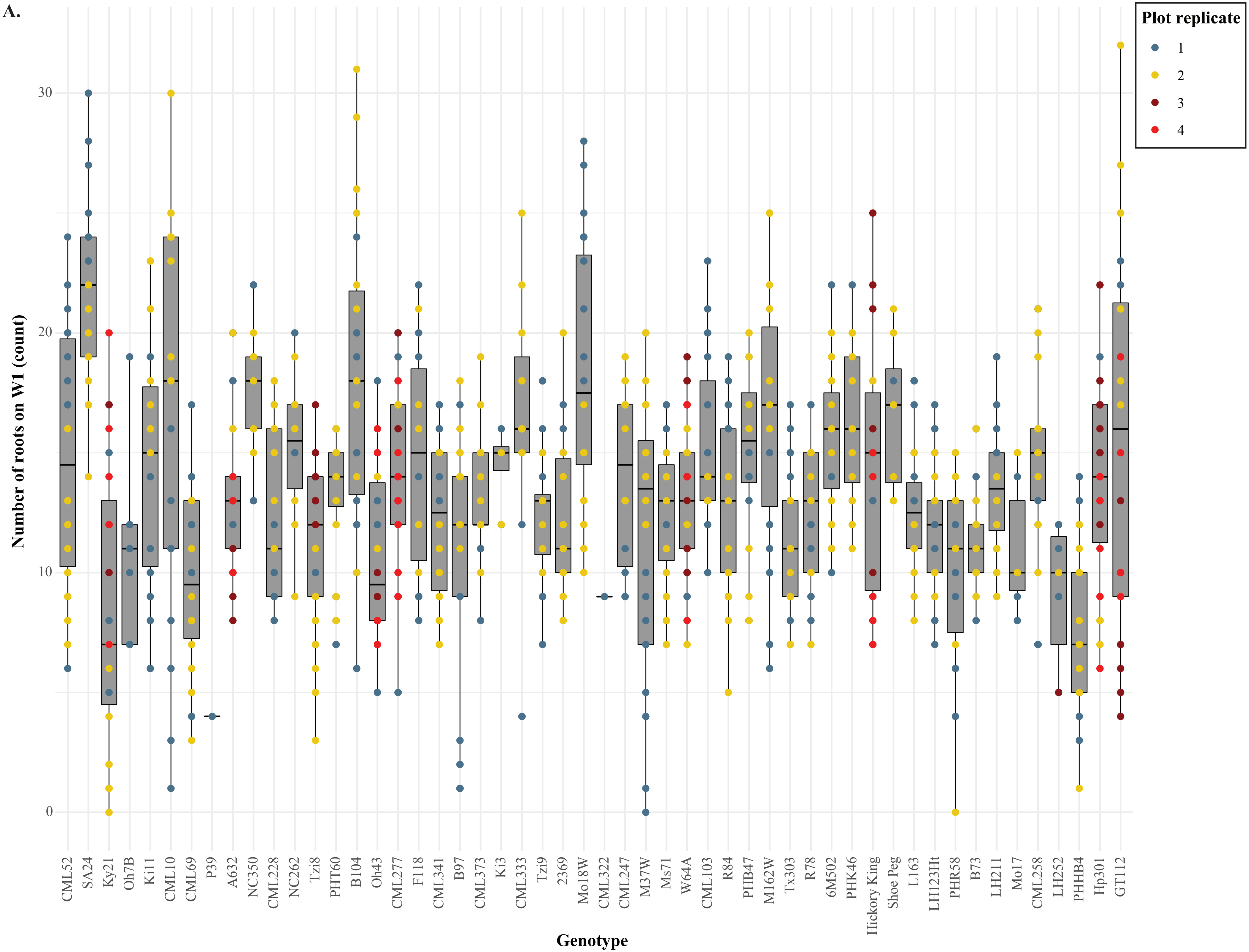

Plot replicate
1
2
3
4
Number of roots on W1 (count)
0
10
20
30
Genotype
CML52
SA24
Ky21
Oh7B
Ki11
CML10
CML69
P39
A632
NC350
CML228
NC262
Tzi8
PHT60
B104
Oh43
CML277
F118
CML341
B97
CML373
Ki3
CML333
Tzi9
2369
Mo18W
CML322
CML247
M37W
Ms71
W64A
CML103
R84
PHB47
M162W
Tx303
R78
6M502
PHK46
Hickory King
Shoe Peg
L163
LH123Ht
PHR58
B73
LH211
Mo17
CML258
LH252
PHHB4
Hp301
GT112

B.

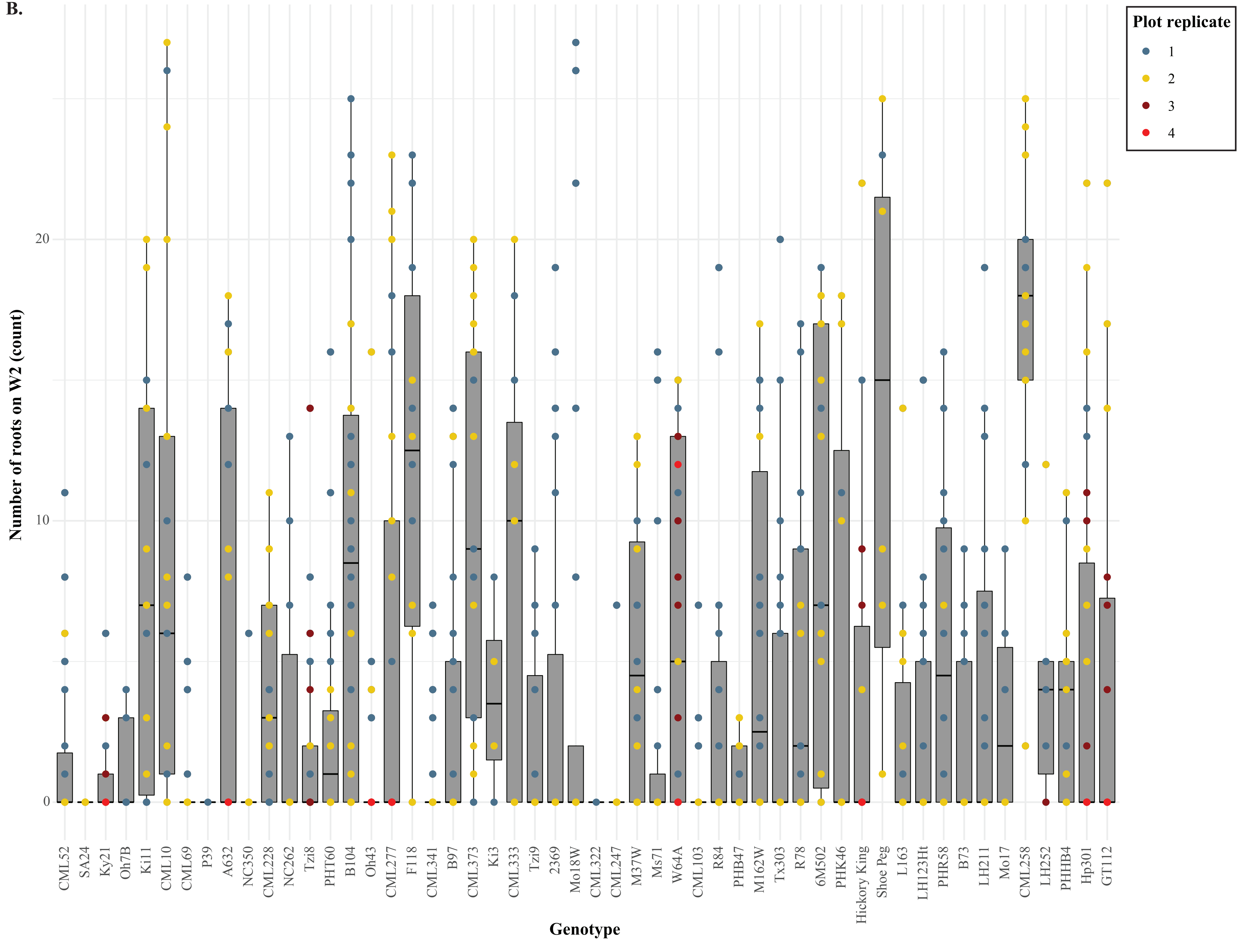
Plot replicate
1
2
3
4
Number of roots on W2 (count)
0
10
20
Genotype
CML52
SA24
Ky21
Oh7B
Ki11
CML10
CML69
P39
A632
NC350
CML228
NC262
Tzi8
PHT60
B104
Oh43
CML277
F118
CML341
B97
CML373
Ki3
CML333
Tzi9
2369
Mo18W
CML322
CML247
M37W
Ms71
W64A
CML103
R84
PHB47
M162W
Tx303
R78
6M502
PHK46
Hickory King
Shoe Peg
L163
LH123Ht
PHR58
B73
LH211
Mo17
CML258
LH252
PHHB4
Hp301
GT112

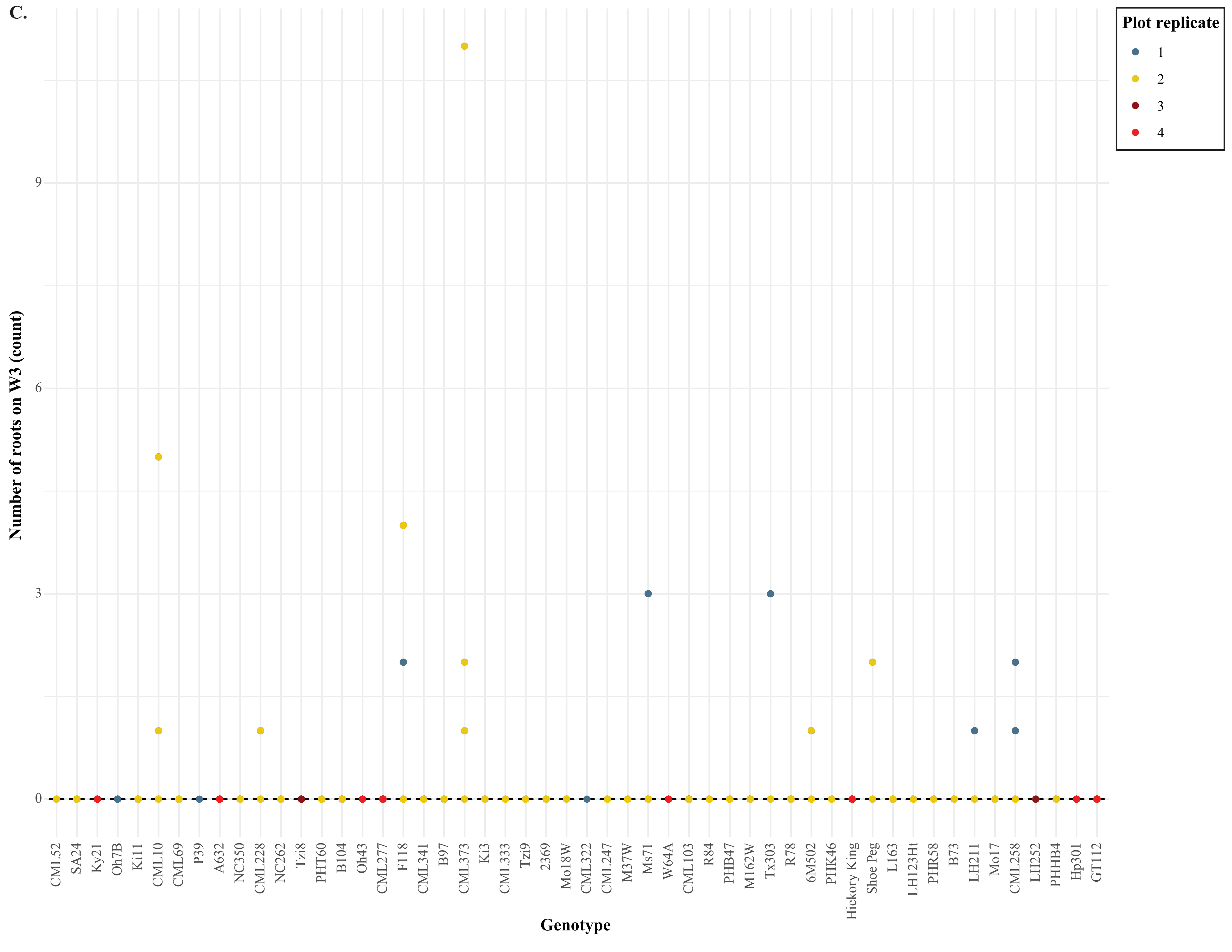
C.
Plot replicate
1
2
3
4
Number of roots on W3 (count)
0
3
6
9
Genotype
CML52
SA24
Ky21
Oh7B
Ki11
CML10
CML69
P39
A632
NC350
CML228
NC262
Tzi8
PHT60
B104
Oh43
CML277
F118
CML341
B97
CML373
Ki3
CML333
Tzi9
2369
Mo18W
CML322
CML247
M37W
Ms71
W64A
CML103
R84
PHB47
M162W
Tx303
R78
6M502
PHK46
Hickory King
Shoe Peg
L163
LH123Ht
PHR58
B73
LH211
Mo17
CML258
LH252
PHHB4
Hp301
GT112

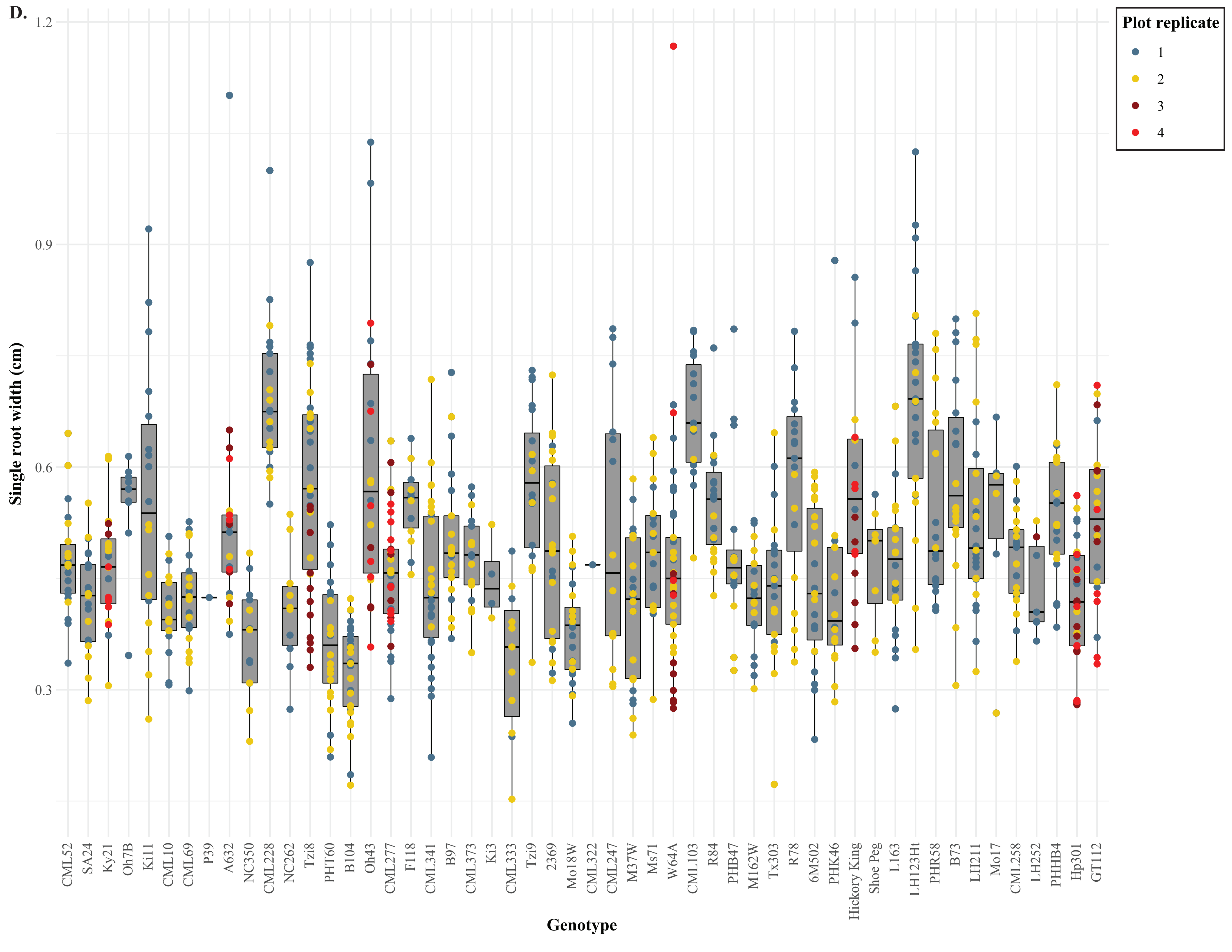

D.
Single root width (cm)
1.2
0.9
0.6
0.3
Genotype
CML52
SA24
Ky21
Oh7B
Ki11
CML10
CML69
P39
A632
NC350
CML228
NC262
Tzi8
PHT60
B104
Oh43
CML277
F118
CML341
B97
CML373
Ki3
CML333
Tzi9
2369
Mo18W
CML322
CML247
M37W
Ms71
W64A
CML103
R84
PHB47
M162W
Tx303
R78
6M502
PHK46
Hickory King
Shoe Peg
L163
LH123Ht
PHR58
B73
LH211
Mo17
CML258
LH252
PHHB4
Hp301
GT112
Plot replicate
1
2
3
4

E.

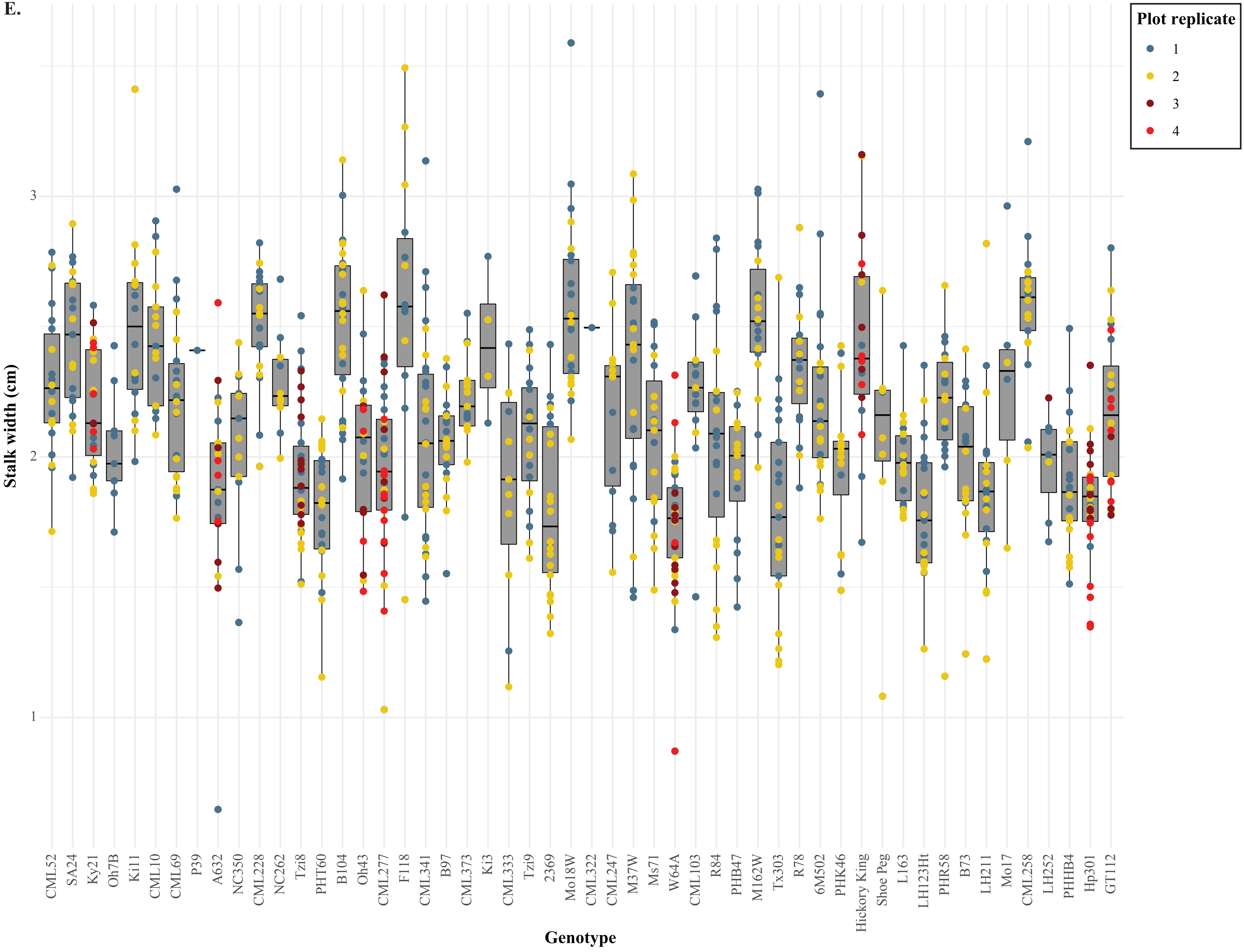

Plot replicate
1
2
3
4
Stalk width (cm)
3
2
1
CML52
SA24
Ky21
Oh7B
Ki11
CML10
CML69
P39
A632
NC350
CML228
NC262
Tzi8
PHT60
B104
Oh43
CML277
F118
CML341
B97
CML373
Ki3
CML333
Tzi9
2369
Mo18W
CML322
CML247
M37W
Ms71
W64A
CML103
R84
PHB47
M162W
Tx303
R78
6M502
PHK46
Hickory King
Shoe Peg
L163
LH123Ht
PHR58
B73
LH211
Mo17
CML258
LH252
PHHB4
Hp301
GT112
Genotype

F.

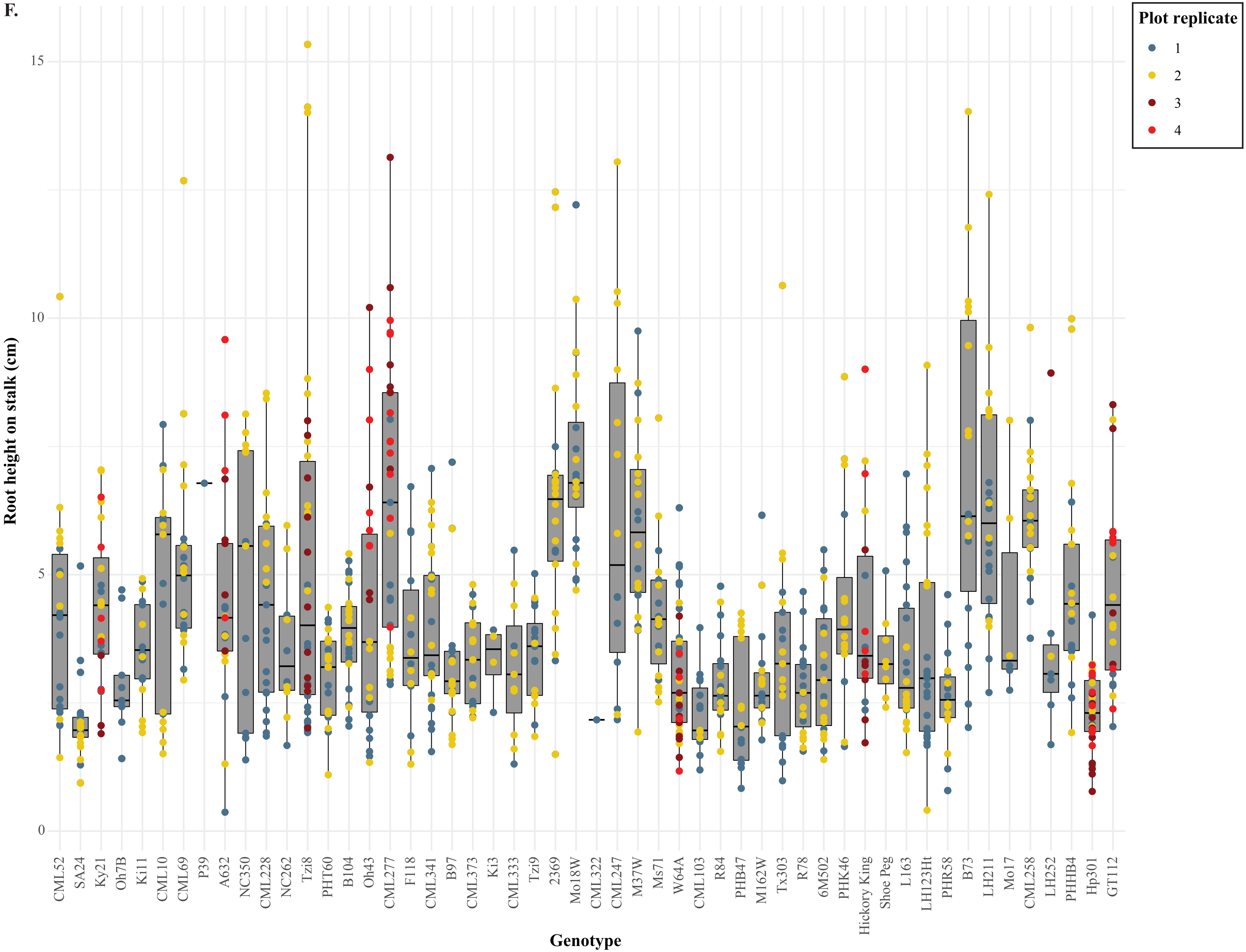
Plot replicate
1
2
3
4
Root height on stalk (cm)
0
5
10
15
Genotype
CML52
SA24
Ky21
Oh7B
Ki11
CML10
CML69
P39
A632
NC350
CML228
NC262
Tzi8
PHT60
B104
Oh43
CML277
F118
CML341
B97
CML373
Ki3
CML333
Tzi9
2369
Mo18W
CML322
CML247
M37W
Ms71
W64A
CML103
R84
PHB47
M162W
Tx303
R78
6M502
PHK46
Hickory King
Shoe Peg
L163
LH123Ht
PHR58
B73
LH211
Mo17
CML258
LH252
PHHB4
Hp301
GT112

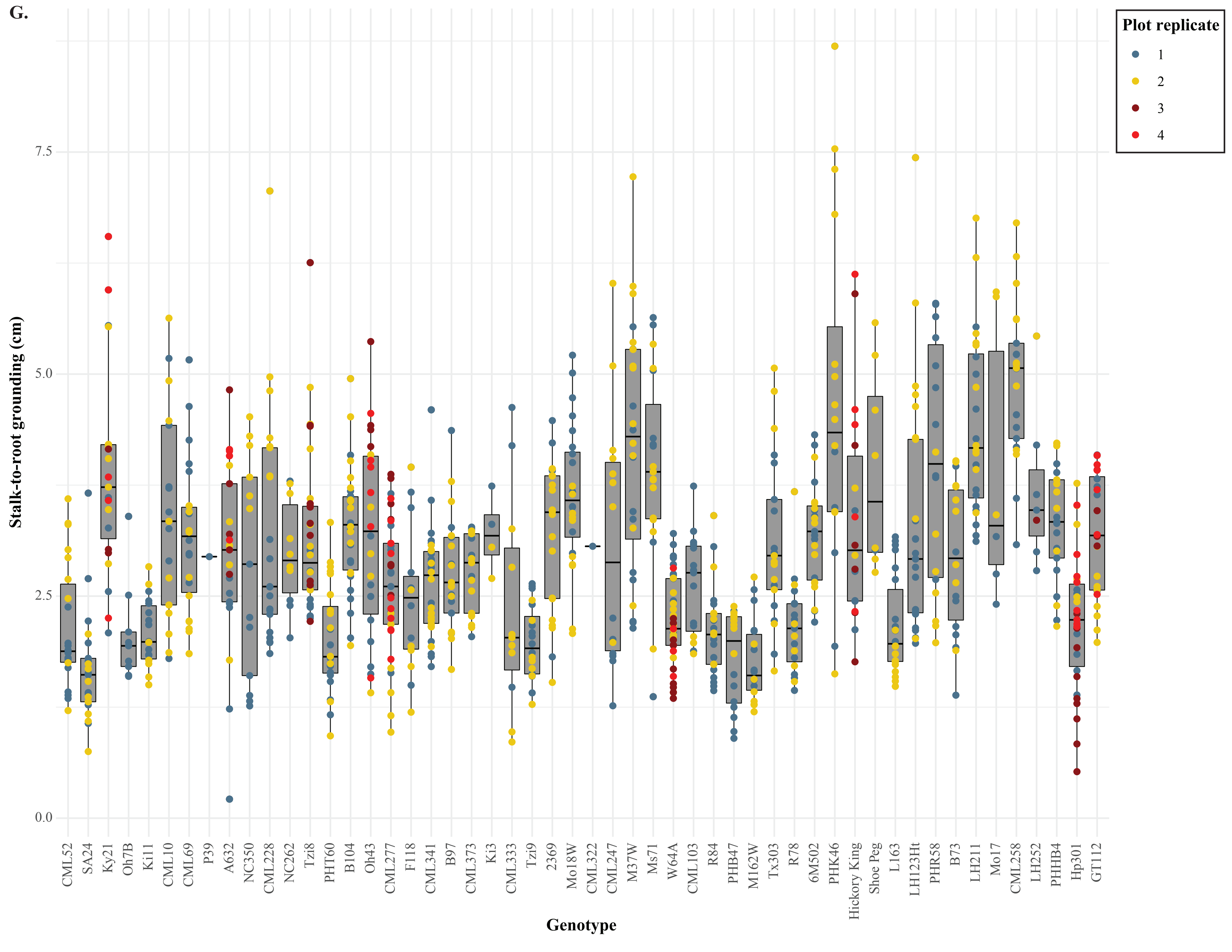

G.
Plot replicate
1
2
3
4
Stalk-to-root grounding (cm)
0.0
2.5
5.0
7.5
CML52
SA24
Ky21
Oh7B
Ki11
CML10
CML69
P39
A632
NC350
CML228
NC262
Tzi8
PHT60
B104
Oh43
CML277
F118
CML341
B97
CML373
Ki3
CML333
Tzi9
2369
Mo18W
CML322
CML247
M37W
Ms71
W64A
CML103
R84
PHB47
M162W
Tx303
R78
6M502
PHK46
Hickory King
Shoe Peg
L163
LH123Ht
PHR58
B73
LH211
Mo17
CML258
LH252
PHHB4
Hp301
GT112
Genotype

H.

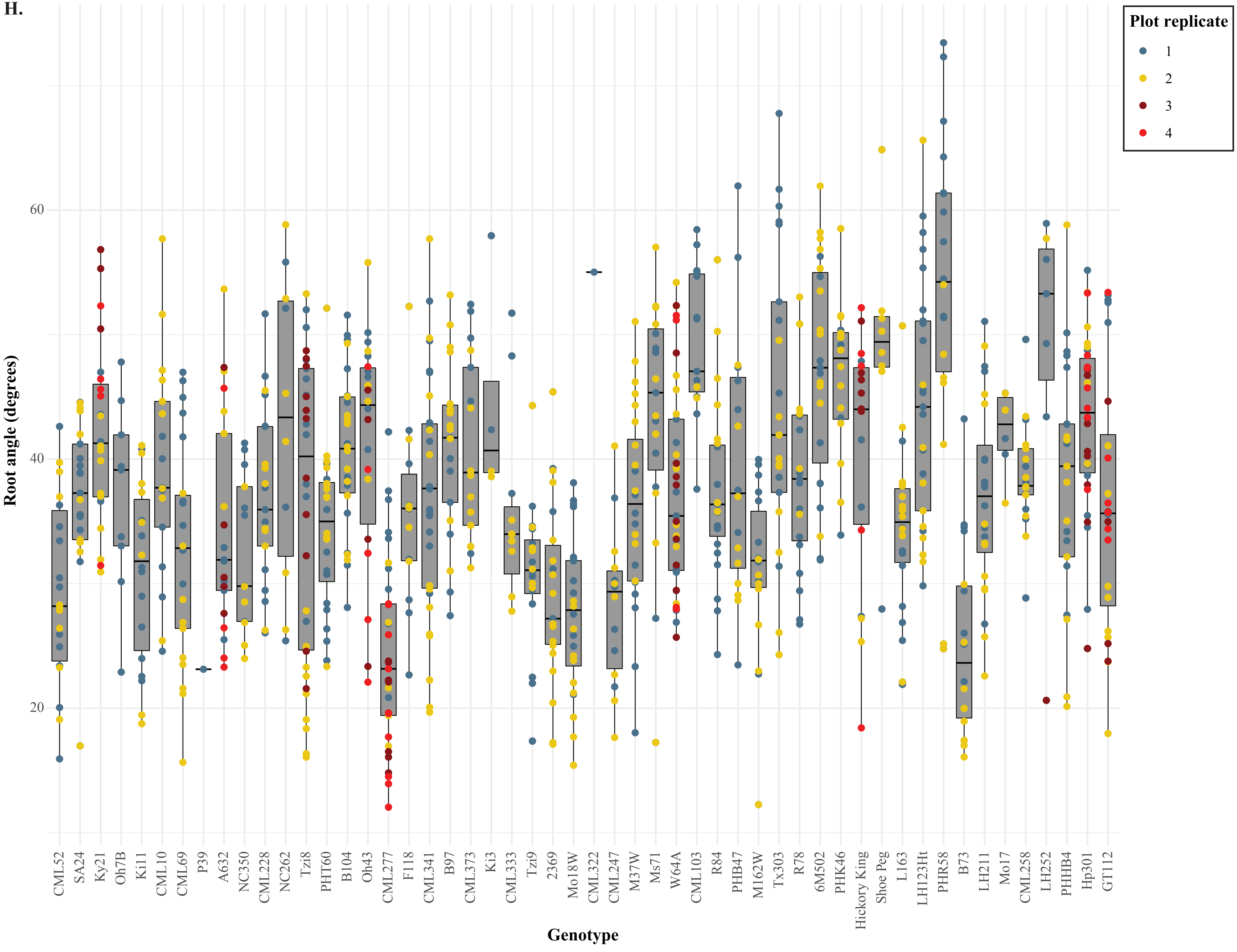
Plot replicate
1
2
3
4
Root angle (degrees)
60
40
20
Genotype
CML52
SA24
Ky21
Oh7B
Ki11
CML10
CML69
P39
A632
NC350
CML228
NC262
Tzi8
PHT60
B104
Oh43
CML277
F118
CML341
B97
CML373
Ki3
CML333
Tzi9
2369
Mo18W
CML322
CML247
M37W
Ms71
W64A
CML103
R84
PHB47
M162W
Tx303
R78
6M502
PHK46
Hickory King
Shoe Peg
L163
LH123Ht
PHR58
B73
LH211
Mo17
CML258
LH252
PHHB4
Hp301
GT112

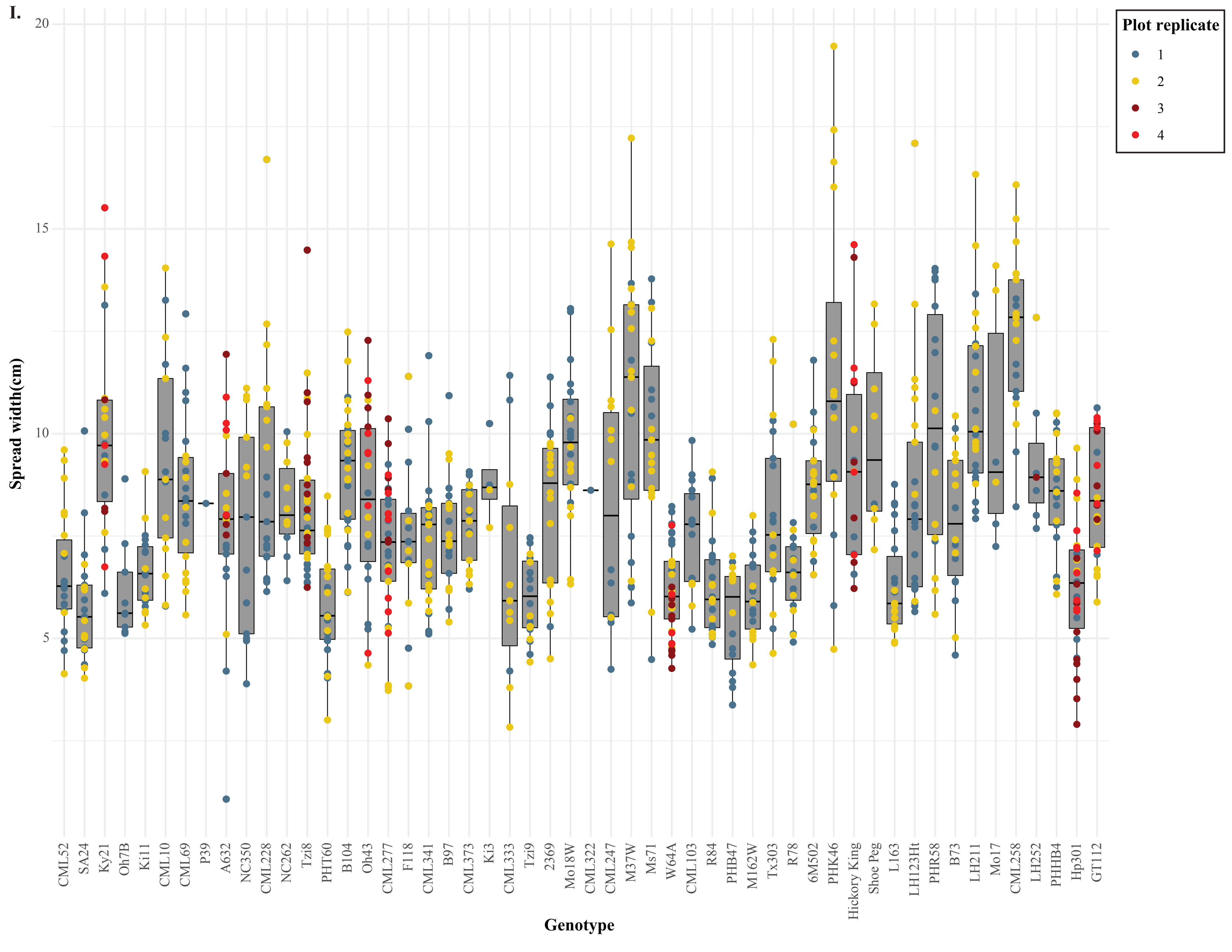

I.
Spread width(cm)
20
15
10
5
Plot replicate
1
2
3
4
CML52
SA24
Ky21
Oh7B
Ki11
CML10
CML69
P39
A632
NC350
CML228
NC262
Tzi8
PHT60
B104
Oh43
CML277
F118
CML341
B97
CML373
Ki3
CML333
Tzi9
2369
Mo18W
CML322
CML247
M37W
Ms71
W64A
CML103
R84
PHB47
M162W
Tx303
R78
6M502
PHK46
Hickory King
Shoe Peg
L163
LH123Ht
PHR58
B73
LH211
Mo17
CML258
LH252
PHHB4
Hp301
GT112
Genotype

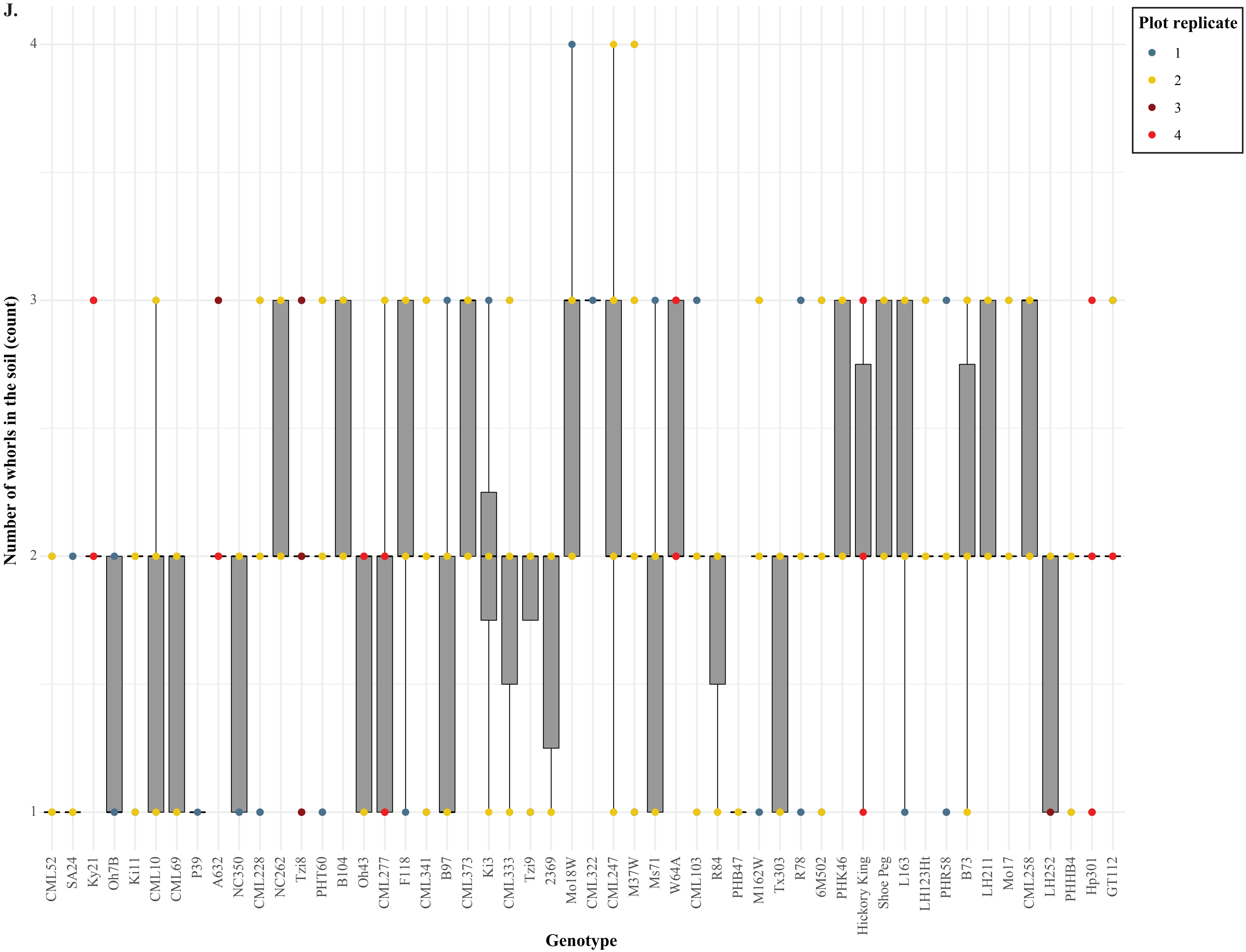
J.
Plot replicate
1
2
3
4
Number of whorls in the soil (count)
1
2
3
4
CML52
SA24
Ky21
Oh7B
Ki11
CML10
CML69
P39
A632
NC350
CML228
NC262
Tzi8
PHT60
B104
Oh43
CML277
F118
CML341
B97
CML373
Ki3
CML333
Tzi9
2369
Mo18W
CML322
CML247
M37W
Ms71
W64A
CML103
R84
PHB47
M162W
Tx303
R78
6M502
PHK46
Hickory King
Shoe Peg
L163
LH123Ht
PHR58
B73
LH211
Mo17
CML258
LH252
PHHB4
Hp301
GT112
Genotype

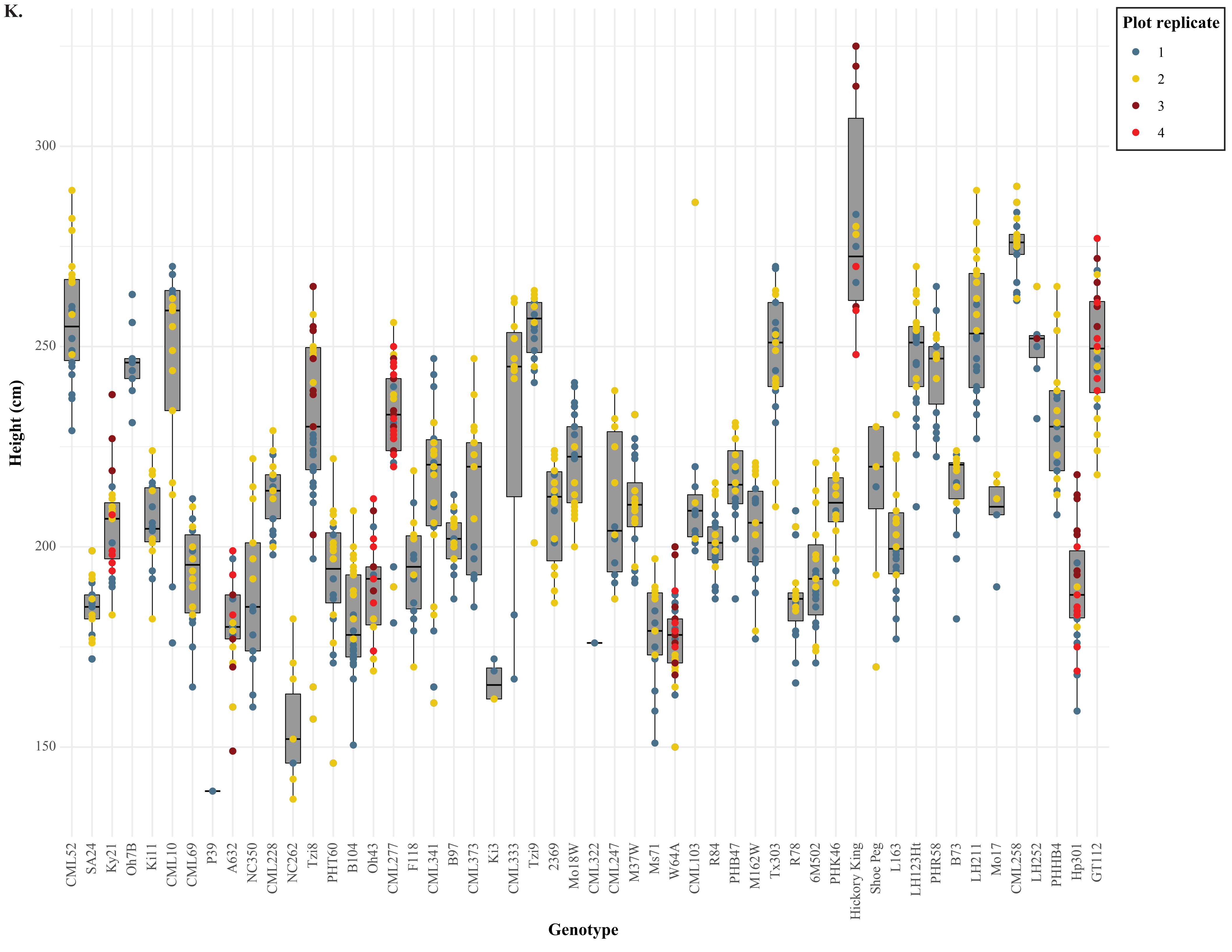
K.
Plot replicate
1
2
3
4
Height (cm)
300
250
200
150
Genotype
CML52
SA24
Ky21
Oh7B
Ki11
CML10
CML69
P39
A632
NC350
CML228
NC262
Tzi8
PHT60
B104
Oh43
CML277
F118
CML341
B97
CML373
Ki3
CML333
Tzi9
2369
Mo18W
CML322
CML247
M37W
Ms71
W64A
CML103
R84
PHB47
M162W
Tx303
R78
6M502
PHK46
Hickory King
Shoe Peg
L163
LH123Ht
PHR58
B73
LH211
Mo17
CML258
LH252
PHHB4
Hp301
GT112

**Figure S5. Plant phenotypes vary among genotypes.** The following phenotypes vary among genotypes: (A) The number of roots on whorl 1 (the whorl closest to the ground, bottom whorl), (B) the number of roots on whorl 2 (middle whorl), (C) the number of roots on whorl 3 (top whorl), (D) the single root width, (E) the stalk width, (F) the root height on stalk, (G) the stalk-to-root grounding, (H) the root angle, (I) the spread width, (J) the number of whorls in the ground, and (K) plant height. (A-K) Genotypes are ordered by rank (high to low) for the None/All ratio according to Figure 1. The color of each dot illustrates the replicate plot where the phenotype data is from. Outliers are outlined in black.

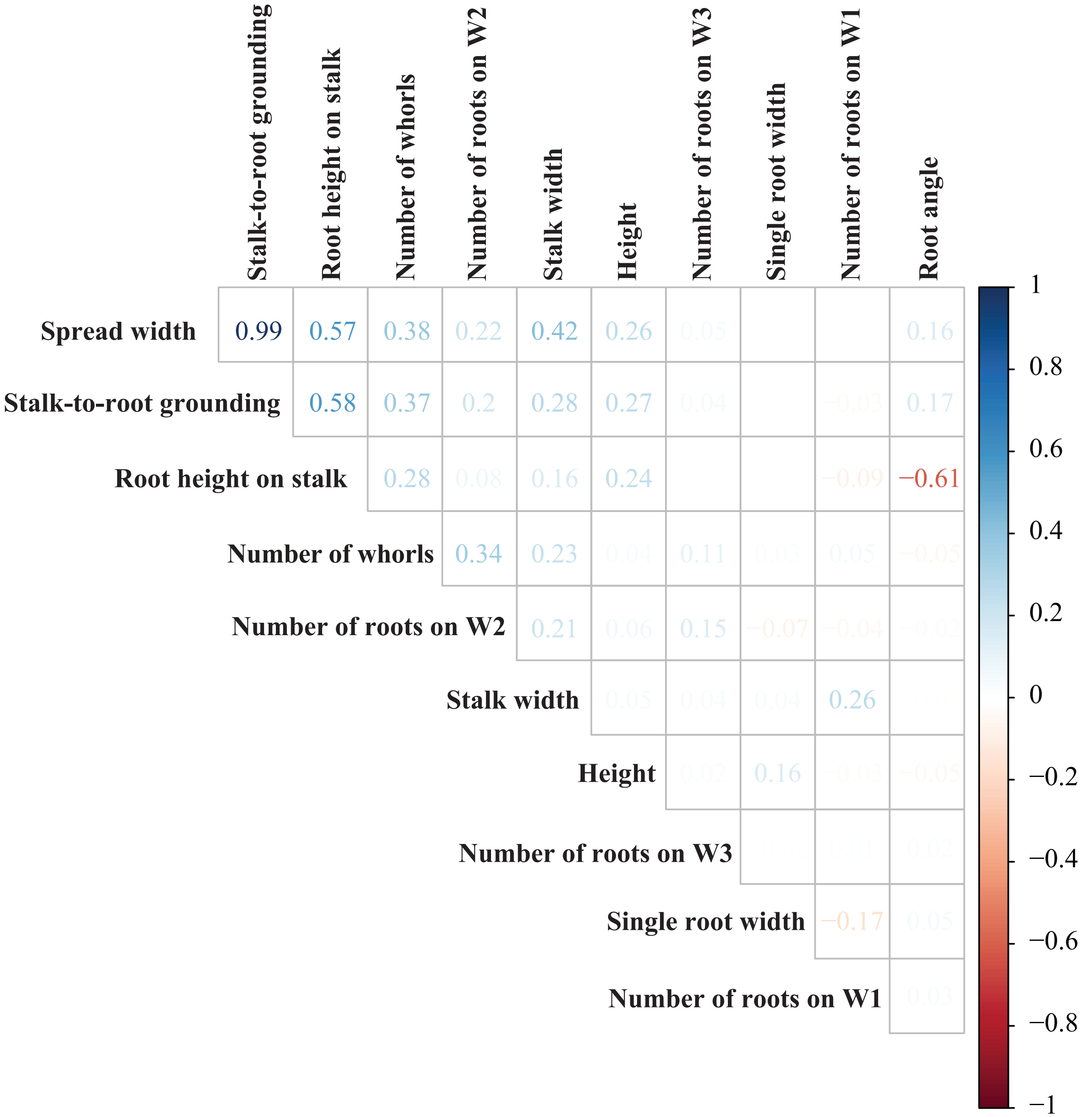


**Figure S6. Brace root phenotypes are correlated with each other.** A Pearson correlation analysis shows which phenotypes are highly correlated.

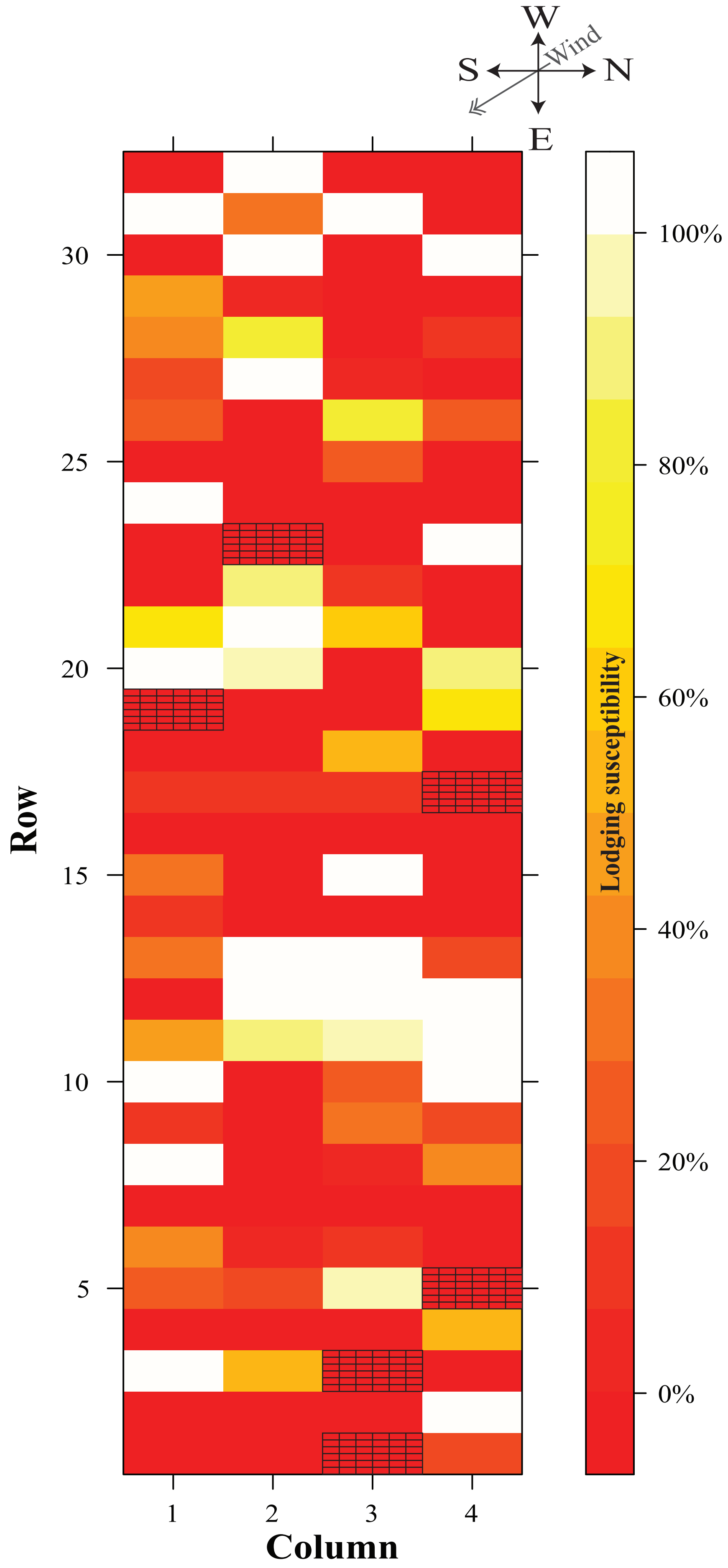


**Figure S7. Root lodging was not related to field position.** Winds from the West and North-West induced root lodging. Plots were colored according to the susceptibility of root lodging within each plot. Plots with 100% root lodging were highlighted with white, whereas plots that had 0% root lodging were highlighted with dark red. Plots with a grid overlaid indicate plots that did not germinate.

**Literature Cited**


Flint-Garcia, S. A., Thuillet, A.-C., Yu, J., Pressoir, G., Romero, S. M., Mitchell, S. E., Doebley, J., Kresovich, S., Goodman, M. M., & Buckler, E. S. 2005. Maize association population: a high-resolution platform for quantitative trait locus dissection. *The Plant Journal*, *44*(6), 1054–1064. https://doi.org/10.1111/j.1365-313X.2005.02591.x

Liu, K., Goodman, M., Muse, S., Smith, J. S., Buckler, E., & Doebley, J. 2003. Genetic structure and diversity among maize inbred lines as inferred from DNA microsatellites. *Genetics*, *165*(4), 2117–2128. https://doi.org/10.1093/genetics/165.4.2117
